# Recurrent impairments in visual perception and place avoidance across autism models are causally linked in the haploinsufficiency model of intellectual disability *Setd5*

**DOI:** 10.1101/2022.10.11.511691

**Authors:** Laura E. Burnett, Peter Koppensteiner, Olga Symonova, Tomás Masson, Tomas Vega-Zuniga, Ximena Contreras, Thomas Rülicke, Ryuichi Shigemoto, Gaia Novarino, Maximilian Jösch

## Abstract

Despite the diverse genetic origins of autism spectrum disorders (ASDs), affected individuals share strikingly similar and correlated behavioural traits that include perceptual and sensory processing challenges. Notably, the severity of these sensory symptoms is often predictive of the expression of other autistic traits. However, the origin of these perceptual deficits remains largely elusive. Here, we show a recurrent impairment in visual threat perception that is similarly impaired in three independent models of ASD with different molecular aetiologies. Interestingly, this deficit is associated with reduced avoidance of threatening environments - a non-perceptual trait. Focusing on a common cause of ASDs, the *Setd5* gene mutation, we define the molecular mechanism. We show that the perceptual impairment is caused by a potassium channel (Kv1) mediated hypoexcitability in a subcortical node essential for the initiation of escape responses, the dorsal periaqueductal grey (dPAG). Targeted pharmacological Kv1 blockade rescued both perceptual and place avoidance deficits, causally linking seemingly unrelated trait deficits to the dPAG. Our findings reveal a link between rapid perception controlled by subcortical pathways and appropriate learned interactions with the environment, and define a non-developmental source of such deficits in ASD.

## Introduction

Autism spectrum disorders (ASD) are conditions characterised by challenges with social interactions and repetitive behaviours, reflected in inadequate responses to others’ mental states and emotions[1,2]. These alterations in social cognition co-occur with disordered sensory processing[3], a widespread yet often overlooked feature across ASD observed in every sensory modality[4]. In the visual domain, affected individuals frequently exhibit difficulties with visual attention and hyper-or hyposensitivity to visual stimuli[5], and mounting evidence suggests that the circuits involved in visual information processing are disrupted[6]. For example, atypical neuronal responses to faces and looming stimuli have been observed in individuals with ASD[7–9]. Such sensory deficits directly affect visually-guided behaviours such as gaze control, which emerges at a few months of age[10] and forms a prominent diagnostic feature[11]. Changes in gaze dynamics to subliminal stimuli[12] indicate that neuronal circuits mediating subconscious visual responses are affected, pointing towards subcortical circuits, particularly the superior colliculus (SC)[6,13,14].

Deficits in sensation are seemingly separate phenomena from the more studied cognitive impairments in ASD, believed to arise from cortical malfunctions[15–17]. However, their close association indicates the possibility of a common underlying thread, particularly because their severity is associated with the strength and expression of other ASD traits[18–20]. Such evidence has recently implicated SC pathway impairments with atypical sensory and cognitive processing[6,14]. Given this evidence, we decided to directly explore the relationship between sensation and cognition in ASD within an innate, robust and reproducible sensorimotor transformation known to be mediated by the SC - the looming escape response (LER)[21,22]. Using *Setd5*[23] as a case study, we show that the LER is impaired, not due to direct sensory or motor deficits. While all animals immediately detect the looming stimulus and can respond vigorously to a threat, they require longer to initiate the escape response than their wild-type siblings and do not form an appropriate aversion to the threat area. These two behavioural traits are highly correlated. The stronger the perceptual deficits, the weaker the avoidance. We further show that these behavioural correlations are also present in aetiologically distinct ASD mouse models (*Cul3*[24] and *Ptchd1*[25]), indicating that these various molecular dysfunctions converge on common behavioural deficits. In *Setd5,* these deficits emerge through changes in the intrinsic excitability due to an increased potassium channel conductance of neurons in the dorsal periaqueductal grey (dPAG), a structure known for commanding escape responses and receiving direct input from the SC[26,27]. Rescuing this hypoexcitability phenotype in the dPAG in adult mice recovers both the perceptual and place avoidance deficits, linking a sensorimotor disorder via a defined molecular mechanism to cognitive dysfunction. Our results show that dPAG dysfunctions can be instrumental in the emergence of symptoms associated with ASDs and that some of the dysfunctions are not developmental, opening a path for their targeted treatment.

## Results

### Delayed looming escape responses

To investigate whether subcortical visuomotor transformations could be affected by genetic mutations associated with ASD, we evaluated the behavioural responses to the innate looming escape response (LER) paradigm. LER is largely independent of cortical input, requiring the SC and its downstream targets to initiate appropriate responses. For this purpose, we tested a genetic haploinsufficiency ASD mouse model affecting the *SET-domain containing 5 (Setd5)*[23], a histone-associated protein found in ∼1 % of patients with intellectual disability and ASD[23]. We subjected mutants (i.e., *Setd5^+/-^*) and their wild-type (WT) siblings to our LER paradigm, where mice would automatically trigger a sequence of five consecutive looms upon entering a threat zone (Fig. 1a, b, see methods). For all experiments, we only used animals without any craniofacial abnormalities, tooth displacement, or eye abnormalities, which are described to sometimes occur in our *Setd5* mutant animals[23]. As previously reported, *Setd5^+/-^* animals were, on average, slightly smaller than their WT siblings[23]. Please note that other models for the same gene show more drastic abnormalities not observed in our hands[28]. During the acclimatisation period, all animals showed similar exploratory behaviour and dynamics (Fig. S1a-g), and seemingly similar behavioural responses, escaping to the shelter upon stimulus presentation (Video S1). However, behavioural differences between the genotypes became evident when observing their reaction times (Fig. 1c). While WT siblings responded robustly to the first stimulus presentation, their heterozygous siblings required, on average, more repetitions to elicit an escape response (Fig. 1d), triggered more looms (Fig. 1e), made more shelter exits (Fig. 1f), had longer reaction times and escaped with less vigour, reaching lower maximum speeds[29] (Fig. 1g). The differences across genotypes were independent of the location, speed of the animal at stimulus onset or heading angle (Fig. S1h-j), or sex (Fig. 1c-j - females, and Fig. S1l-n-males). The difference in response reaction time is an intrinsic and not an adaptive property, since it is present from the first loom presentation the animal encounters (Fig. 1hg, left). However, the escape vigour of the *Setd5*^+/-^ animals during these first-ever encountered loom presentations was indistinguishable from their WT siblings (Fig. 1h, right). This indicates that the escape delay differences are not due to intrinsic motor problems, but emerge due to perceptual decision-making deficits - the process of extracting and integrating sensory information to initiate a subsequent action. These results highlight the LER as a robust behavioural assay to probe perceptual impairments and their underlying circuit dysfunctions in visuomotor processing in ASD.

**Figure 1.**
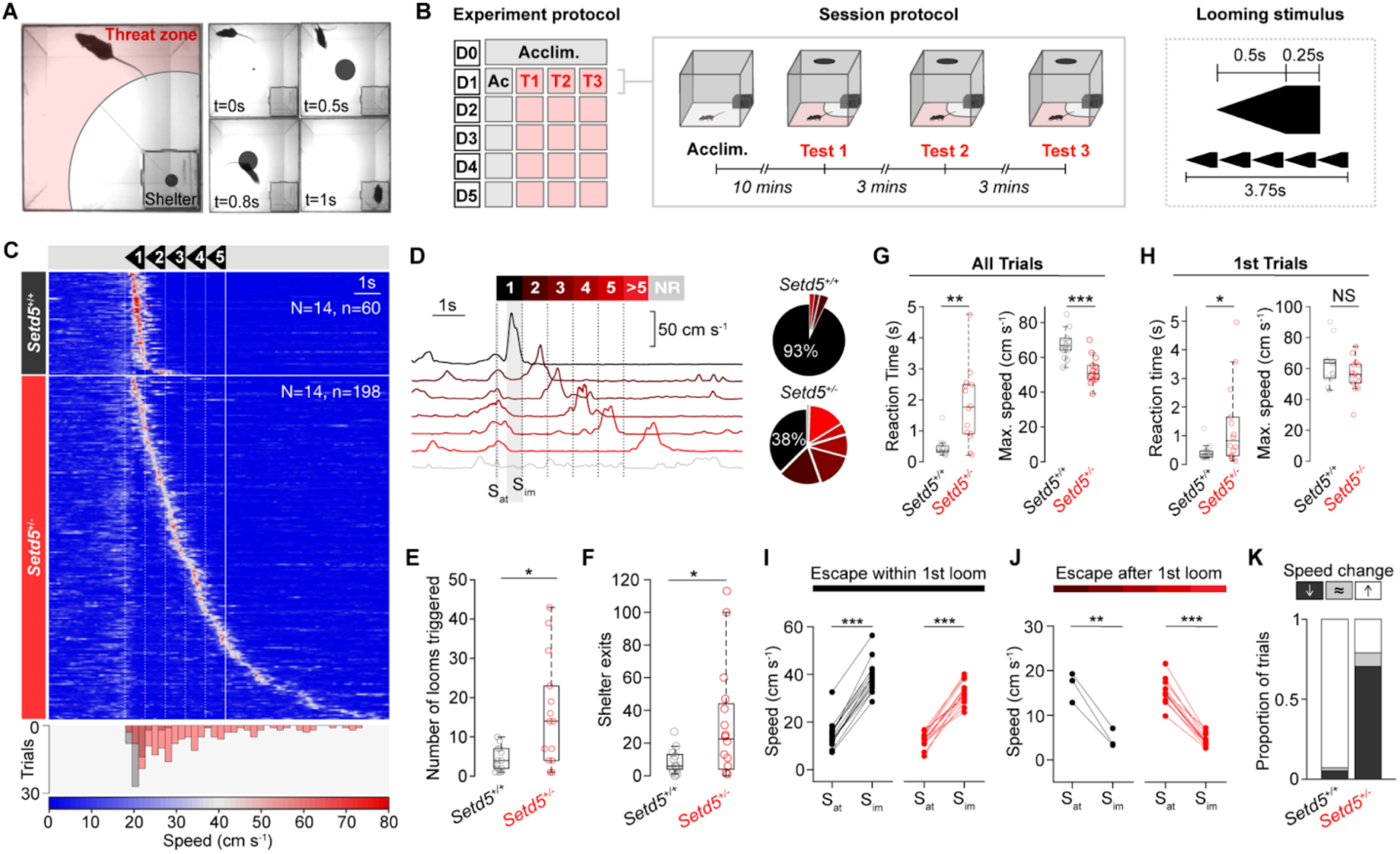
ASD mice exhibit delayed and less vigorous looming escape responses. **a**, Looming escape response (LER) paradigm showing the shelter’s location and the threat zone (left) and LER example (right). **b,** Paradigm schematic. Day 0 (D0) was used for acclimatisation. D1-D5 consisted of an acclimatisation period (grey) followed by three LER tests (red). The looming stimulus consisted of 5 consecutive looms (right). **c**, Raster plot of mouse speed during LER (white, dotted vertical lines denote the start of each loom; solid white line denotes the end of the stimulus) for *Setd5^+/+^* (upper, n =14, 60 trials) and *Setd5^+/-^*(lower, n = 14, 198 trials), sorted by reaction time. Bottom, distribution of reaction times for all *Setd5^+/+^* (black) and *Setd5^+/-^* (red) trials (p < 0.001, two-sample Kolmogorov-Smirnov test). **d**, Left, example trials based on whether the mouse responds within 1 of the 5-loom stimuli, after the 5th (>5), or not at all (NR - no response) from one *Setd5^+/-^* mouse. Grey shaded areas represent the frames used to calculate the speed at stimulus onset (Sat) and the immediate response speed (Sim). Right, proportion of escapes to loom presentations. **e**, Total looms triggered across the 5 test days (*Setd5^+/+^*, 4.36 looms; *Setd5^+/-^*, 16 looms, P = 0.013). **f**, Total shelter exits across the 5 test days (*Setd5^+/+^,* 6.0; *Setd5^+/-^,* 24.5, P = 0.019). **g**, Average reaction time (left) and maximum escape speed (right) per animal (reaction time, P = 0.001; max. escape speed, P < 0.001). **h**, as (**g**), but only for the very first loom presentation (reaction time, P = 0.046; max. escape speed, P = 0.166). **i**, Average immediate speed change following the stimulus presentation for all trials where the mice escape within the first loom presentation (left, *Setd5^+/+^,* n = 14, black, p < 0.001; right, *Setd5^+/-^*, n = 14, red, p < 0.001). Sat is the mean speed of the animal ±50 ms of stimulus onset and Sim is the mean speed of the animal 300-800ms after stimulus onset. **j**, as (**i**) but for trials where the mice escape during or after the second loom (left, *Setd5^+/+^,* n = 3, black, p = 0.007; right, *Setd5^+/-^*, n = 11, red, p < 0.001). **k**, Proportion of response types per genotype (X^2^ = 103.9, p < 0.001, X^2^ test of independence). P-values: Wilcoxon’s test, p-values: paired *t*-test, unless specified.

To test whether the delay in initiating an escape to the loom was due to the mutant mice simply detecting the stimulus later than their WT siblings, we examined the immediate change in the speed of the animals from the time of the first loom, selecting only the trials where the mice performed an escape to the shelter at any point during the looming events. For trials where the animals generated an escape within the first loom presentation, both WT and mutant animals increased their speed significantly upon stimulus onset (Fig. 1i). For trials where the mice didn’t generate an escape within the first loom, which was the majority of trials for mutant animals but only a small fraction of trials for WT animals (Fig. 1d), the mice significantly reduced their speed in the time immediately following the stimulus onset (Fig. 1J). This demonstrates that *Setd5*^+/-^ animals detect and respond to the stimulus within a similar time frame as their WT siblings, but preferentially perform a locomotor arrest instead of an escape response (see Supplementary Videos). This arrest behaviour is more reminiscent of risk assessment behaviour as previously shown for loom stimuli of different contrast[29] rather than the defensive freezing response characterised by the sustained cessation of all movement, since the animals were still performing small movements of their head and upper body. Indicating that the animals are using this time to perform ongoing threat evaluation. This suggests that the delay in LER arises from difficulties in either evaluating the threat level of the stimulus or initiating an appropriate response.

Given that cortical malfunctions have been suggested as the cause of sensory differences in autism[15–17] and that the visual cortex has been shown to modulate the response magnitude of looming sensitive cells in the SC[30], we tested whether the behavioural responses are affected in cortex and hippocampus-specific conditional *Setd5* animals (*Setd5^+/fl^*; *Emx1*-Cre, Fig. S2). These animals showed no behavioural differences in their reaction times and vigour, or response kinetics (Fig. S2c-f), in line with previous studies that show that subcortical pathways[29,31], namely the SC and PAG, and not altered top-down modulation from cortical areas, are required for this behaviour.

### Repetitive looming escape responses

Although the innate LER does not require learning, repeated loom presentations cause adaptive behavioural changes, for example, the emergence of place avoidance of the threat zone[29]. Given that *Setd5^+/^*^-^ mice triggered three times more looming events and shelter exits than their WT siblings (Fig. 1e,f), we explored if altered adaptation to repeated presentations could account for the observed difference in average reaction time and vigour (Fig. 1g). For that purpose, we compared the intrinsic behavioural characteristics and the effect of the LER on the exploration strategies and behavioural adaptations across days (Fig. 2a-d). Before stimulus exposure, both cohorts had similar exploratory strategies (Fig. S1a-g), indicating that their innate exploration strategies and thus, intrinsic levels of anxiety, cannot account for the differences in reaction time and vigour. After stimulus exposure, WT animals showed expected reductions in their exploratory behaviour following the initial exposure to the looming stimulus, making fewer exits than during the pre-stimulus exploration time (Fig. 2b), eliciting fewer looms in total (Fig. 2c) with relatively constant reaction time (Fig. 2d). Strikingly, *Setd5^+/-^* consistently triggered more looms and had no signs of sensitisation upon repeated exposures (Fig. 2a-d). After the initial decrease in exploration following the first stimulus presentation, they showed no further consolidation of place avoidance to the threat zone. The reaction time and vigour of the escape response remained longer and slower across days (Fig. 2d), respectively, indicating that *Setd5^+/-^* do not sensitise as their WT siblings do to the LER paradigm. We next tested if the delayed perceptual decisions (Fig.1) are related to the consolidation of place avoidance, two seemingly independent behaviours. We analysed the total number of shelter exits for each animal and compared them with their average reaction times and vigour (Fig. 2e,f). Surprisingly, the strength of the place avoidance was a strong predictor of the reaction times, but not vigour, suggesting that the timing of the perceptual decision and the formation of place avoidance are intrinsically linked. To assess the limits of response adaptation and to ascertain if the presence of a reward would overcome the fear, we conducted the same behavioural experiment but with the presence of a food reward on the far side of the arena within the threat zone (Fig. 2g-j), and with no inter-stimulus interval restriction. Although WT siblings did make some attempts to retrieve the food reward, they quickly sensitised, rarely leaving the shelter and were unsuccessful in obtaining the food reward. *Setd5^+/-^*, on the other hand, increased their shelter-leaving events, often persisting until reaching the reward (Fig. 2g-i, Video S2). Despite the rapid and repeated exposure, *Setd5^+/-^* mice continued to escape to the shelter upon stimulus presentation, for >10 consecutive presentations (Fig. 2h-j). Their reaction times showed a mild adaptation (Fig. 2j, top), while vigour remained largely constant (Fig. 2j, bottom). These results show that two behavioural traits, LER and place avoidance, strongly correlate, suggesting they share a common neuronal pathway.

**Figure 2.**
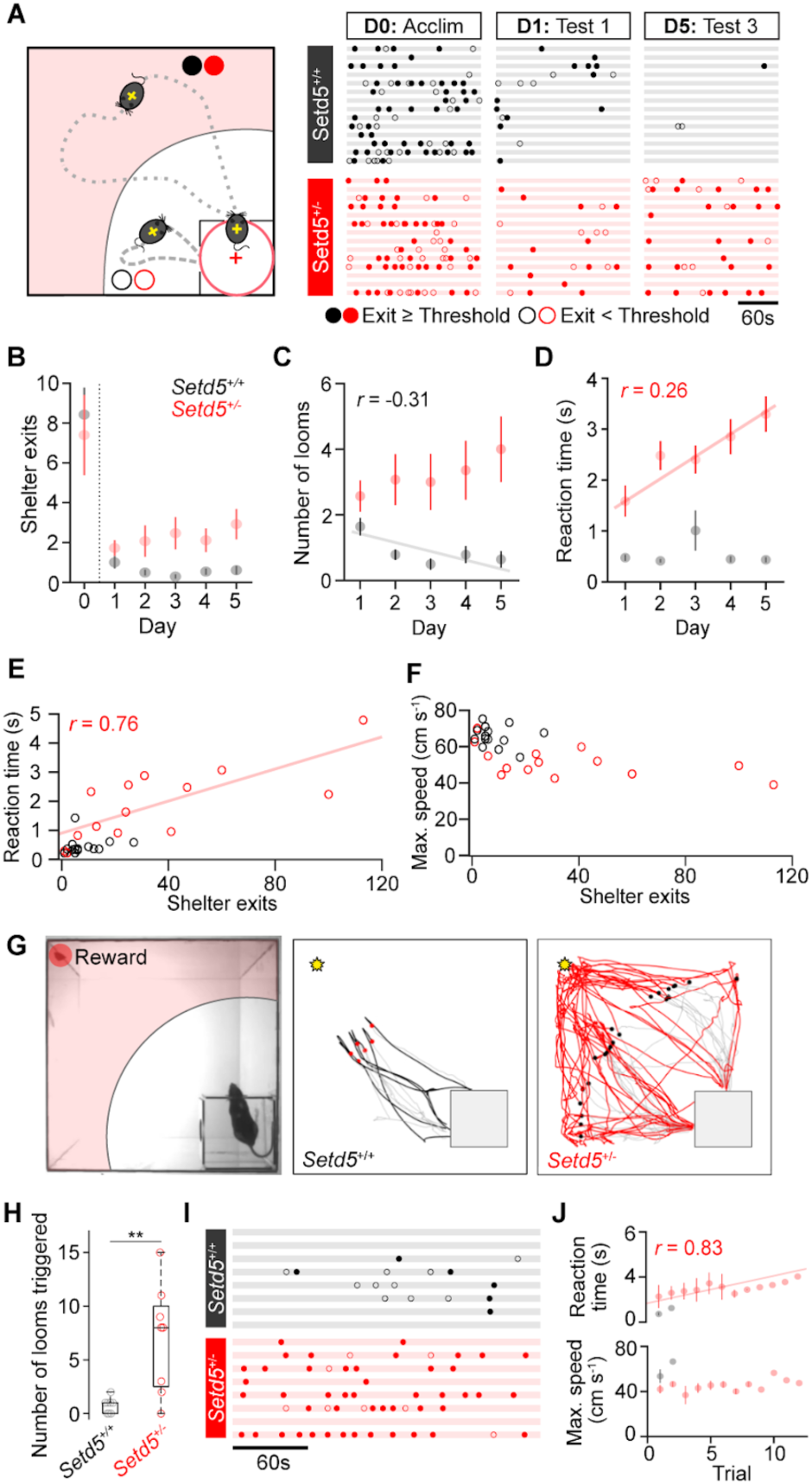
Altered adaptation and repetitive behavioural phenotype to the LER. **a**, left, Graphic depicting exits where the mouse enters (dotted line, filled dot) or does not enter (dashed line, open dot) the threat zone. Right, ethogram of exploratory shelter exit behaviour during the pre-stimulus acclimatisation, the first and last test of the LER paradigm. Each row represents one animal. **b-d,** Adaptation in the number of shelter exits, average number of looms triggered and reaction times across days (**b**, Setd5*^+/+^*: p = 0.2189, *Setd5^+/-^*: p = 0.4974; **c**, *Setd5^+/+^*, p = 0.0087; Setd5*^+/-^*, p = 0.2192; **d**, *Setd5^+/+^*, p = 0.890; Setd5*^+/-^*, p < 0.001). **e**, Relationship between the number of shelter exits and the average reaction time per animal (*Setd5^+/+^,* p = 0.626; *Setd5^+/-^*, p = 0.002). **f**, as (**e**) but for maximum escape speed per animal (*Setd5^+/+^*, p = 0.547; *Setd5^+/-^*, p = 0.053). **g**, Left, reward trial example showing the location of the food reward within the threat zone. Right, example trajectories during the reward trials show the mouse’s position for the 3 s before triggering the loom (light grey) and the 6 s following the stimulus start (black or red). Filled dots represent the position of the mouse when the stimulus was triggered, grey square represents the shelter, and the yellow star shows the position of the food reward. **h**, Number of looms triggered during the reward trial (*Setd5^+/+^*, 1 bout; *Setd5^+/-^*, 8 bouts, P = 0.005). **i**, Ethograms of exits, as in (**a**), during the reward trials show an increased probability of *Setd5*^+/-^ mice leaving the shelter during a trial (number of exits, 2.12 for *Setd5^+/+^,* 6.88 for *Setd5^+/-^*, p = 0.057). **j**, Reaction time (top panel, *Setd5*^+/-^, 44 trials, *r* = 0.828, *p* = 0.0001) and escape vigour (bottom panel, *Setd5*^+/-^, 44 trials, *r* = 0.184, *p* = 0.5116) during repeated presentations of the loom. Trials when the animal was interacting with the reward were excluded. P-values: Wilcoxon’s test, p-values: Pearson’s correlation test, unless specified. Plotted linear fits depict the statistically significant correlations.

#### Convergent behavioural deficits across ASD models

To test whether the deficiencies observed are specific to *Setd5^+/-^*animals or might reveal a more general behavioural phenomenon across ASD models with distinct molecular and genetic aetiologies, we decided to test two additional highly penetrant mutations, *Cullin3* (*Cul3*), a ubiquitin ligase-encoding gene, and *Ptchd1* (Fig. S3a-c), a member of the Patched family speculated to function as a Sonic hedgehog receptor[25]. We exposed sex-matched sibling pairs from these models to the same LER protocol as used for the *Setd5* model and observed qualitatively similar behavioural deficits in reaction time (*Ptchd1*: Fig. 3a-d; *Cul3*: Fig. 3I-l) and place avoidance (*Ptchd1*: Fig. 3e-g; *Cul3*: Fig. 3i-l). Both cohorts required, on average, more repetitions to elicit an escape response (Fig. 3a,b & i,j), had longer reaction times between the stimulus start and the maximum speed (Fig. 3c,k), displayed slower maximum speed compared to their WT siblings (Fig. 3d,l), triggering more looms (Fig. 3e,m) and leaving the shelter more often (Fig. 3f,n), but had no differences in their behavioural strategies prior to the first loom presentation (Fig. 3g, o, Fig. S3d,k). Similar to *Setd5*, these changes were not due to the animals not reacting to the first stimulus, as seen in their consistent arrest behaviour (Fig. S3e,l), nor to differences in their exploratory behaviour (Fig. S3d,k). This difference in response reaction time is an intrinsic and not an adaptive property since it is present during the first loom presentation (Fig. S3g,n). However, they varied in strength, with *Cul3^+/-^*mice showing a less severe phenotype than the *Setd5^+/-^* or *Ptchd1^Y/-^* models. Interestingly, slower reaction times and reduced vigour could also be observed in *Ptchd^Y/-^* animals, despite their baseline hyperactivity (Fig. S3di). Interestingly, in both cohorts, the strength of the place avoidance and the reaction times were significantly lower and longer, respectively (Fig. 3c,f,k,n), being linearly correlated in *Ptchd1^Y/-^*, but not *Cul3^+/-^* (Fig. 3h,p). These results suggest a general link between the timing of the perceptual decision and the formation of place avoidance, despite different molecular and developmental origins.

**Fig. 3.**
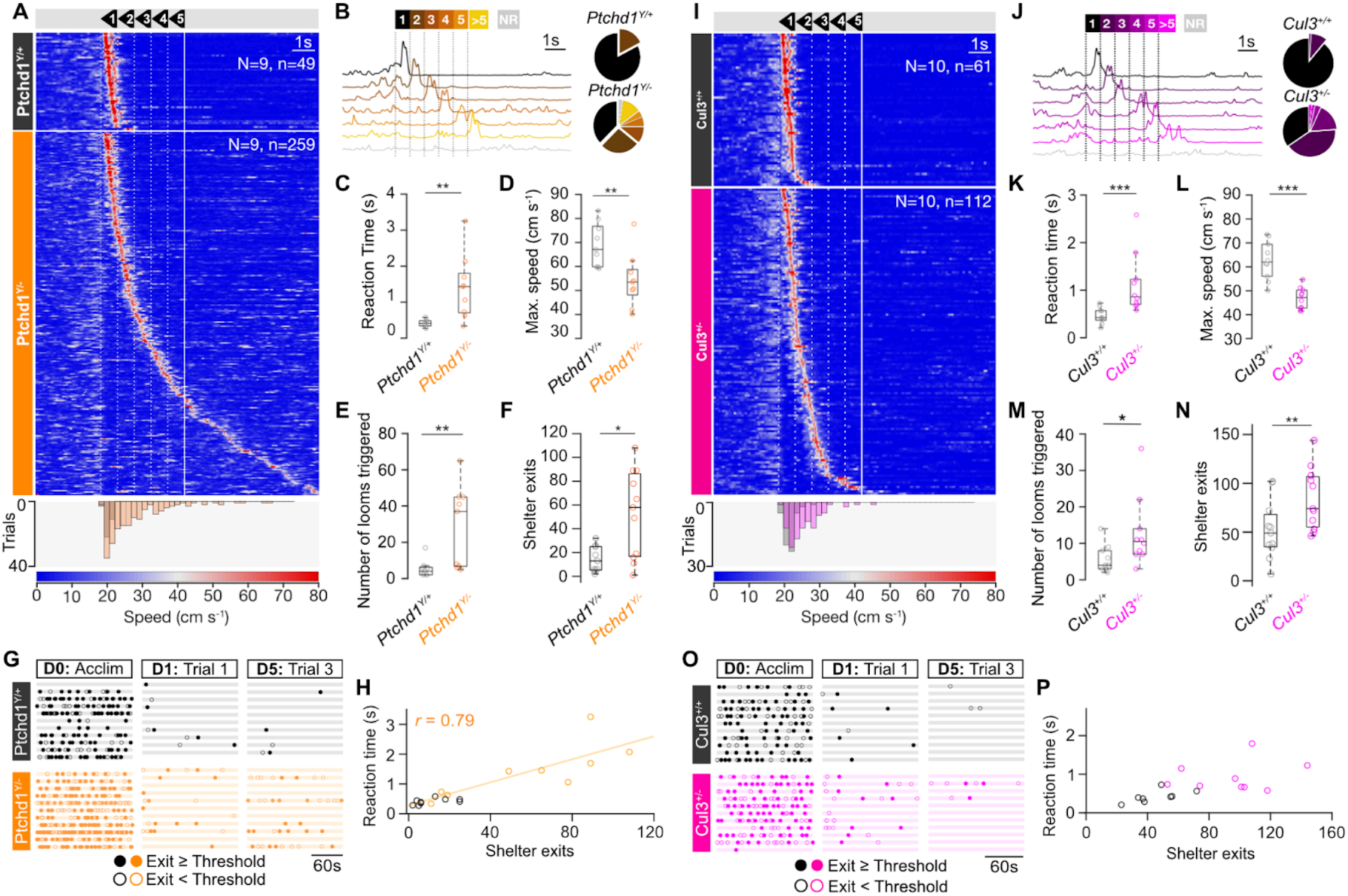
Conserved behavioural changes across etiologically distinct autism models. **a**, Raster plot of mouse speed in response to the looming stimuli for *Ptchd1^Y/+^*(upper, n = 9, 49 trials) and *Ptchd1^Y/-^* (lower, n = 9, 259 trials), sorted by reaction time. Bottom, distribution of reaction times for all *Ptchd1^Y/+^*(black) and *Ptchd1^Y/-^* (orange) trials. **b**, Left, Left, example trials based on whether the mouse responds within 1 of the 5-loom stimuli, after the 5th (>5), or not at all (NR - no response) from one *Ptchd1^Y/-^*mouse. Right, proportion of escape to loom presentations. Trials where mice escaped within the first loom: *Ptchd1^Y/+^*, 0.826, *Ptchd1^Y/-^*, 0.374, p = 0.004. **c**, Average reaction time and (**d**) maximum escape speed per animal, for all trials where the mice escaped (Reaction time; *Ptchd1^Y/+^*, 0.411s, *Ptchd1^Y/-^*, 1.41 s, p = 0.002. Maximum escape speed; *Ptchd1^Y/+^*, 69.1 cms^-1^, *Ptchd1^Y/-^*, 54.2 cms^-1^, p = 0.005). **e**, Average total looms triggered per genotype across 5 days of testing (*Ptchd1^Y/+^*, 5.44 looms; *Ptchd1^Y/-^*, 28.8 looms, P = 0.004). **f**, Average total shelter exits across the 5 test days (*Ptchd1^Y/+^,* 15; *Ptchd1^Y/-^,* 53, P = 0.025). **g**, Ethogram of shelter exits during the pre-stimulus acclimatisation, first and last trial of the LER paradigm. Each row represents one animal, filled and open dots represent exits that crossed into the threat zone or not, respectively. **h**, Relationship between the number of shelter exits and the average reaction time per animal (*Ptchd1*^Y/+^, p = 0.126; *Ptchd1*^Y/-^, *r* = 0.76, p = 0.012, Pearson’s correlation). **i-p**, same as **a-h** but for *Cul3.* **i,** *Cul3*^+/+^ (top, n = 10, 61 trials) and *Cul3*^+/-^ (bottom, n = 10, 112 trials. Bottom, distribution of reaction times (p < 0.001, two-way KS test). **j**, *Cul3*^+/+^, 0.892, *Cul3*^+/-^, 0.347, p < 0.001, two-way KS test). **k**, Reaction time; *Cul3*^+/+^, 0.464 s, *Cul3*^+/-^, 1.11 s, P < 0.001. **i,** Maximum escape speed; *Cul3*^+/+^, 62.7 cm s^-1^, *Cul3*^+/-^, 46.8 cm s^-1^, P < 0.001). **m**, (*Cul3*^+/+^, 5.60 looms; *Cul3*^+/-^, 12.9 looms, P = 0.023). **n**, (*Cul3*^+/+^, 47; *Cul3*^+/-^, 89, P = 0.006). **p**, (*Cul3*^+/+^, p = 0.078; *Cul3*^+/-^, p = 0.571, Pearson’s correlation). Box-and-whisker plots show median, IQR and range. Shaded areas represent SEM. Lines are shaded areas, mean ± s.e.m., respectively. P-values are Wilcoxon’s test, p-values: two-sample Kolmogorov-Smirnov test, unless specified.

### Deep medial SC, not retinal drive, is linked to delay in LER

Changes in average reaction times have previously been shown to depend on the saliency of the threat stimulus. Whereas repeated high-contrast looming stimuli evoke strong LER, low-contrast looms drive less vigorous escapes and longer reaction times[29], suggesting that deficits in visual processing could underlie the behavioural differences observed in *Setd5^+/-^*. This possibility is aggravated by the fact that the expression of genes involved in eye development are known to be misregulated in *Setd5^+/-^* animals[23]. To explore possible visual response changes, we recorded visually evoked responses across different layers of the SC, using 32-channel silicon probes in head-fixed, awake-behaving animals (Fig. S4a), and determined the recording depth using current source density analysis (Fig. S4b-c). In our setup, mutant and control animals had similar running probabilities and showed similar baseline firing properties (Fig. S4d). We next assessed visual response properties to full-field flashes, visual stimuli that drive all neurons similarly, irrespective of their receptive field positions. First, we show that pupil dynamics are unaffected across genotypes (Fig.S 4e). Next, when clustering visually responsive cells into 10 clearly defined groups, we could observe similar proportions and identical response properties (Fig. S4f-g). Identical responses were also observed in their spatio-temporal receptive fields and the proportion of ON and OFF-responsive cells (Fig. S4a-g). Direction-selective responses (Fig. S4h,i) were also indistinguishable across groups. Finally, we explored the response properties to looming stimuli across SC layers. Looms elicit robust and reproducible firing across genotypes (Fig. 4j), with similar firing, peak rates and time-to-peak responses across genotypes (Fig. S4k-m). The overall response kinetics (Fig. S4n) and the adaptation in average firing rates across presentations (Fig. 4o,p) were indistinguishable across genotypes. The lack of any reportable visual response difference across the SC layers indicates that retino-colliculuar visual processing is largely unaltered and excludes the possibility that sensory processing can account for the observed visuomotor deficits and suggests that the main cause of the behavioural difference may reside in downstream areas. Given that optogenetic stimulation of deep medial SC (dmSC) neurons that project to the dorsal periaqueductal grey (dPAG) generates robust escape responses and that the dPAG encodes for the choice of escape and controls the escape vigour[29], we hypothesised that activating dmSC neurons would reveal behavioural differences between genotypes and delineate the underlying circuit impairments. To test this, we focused on the *Setd5* mouse line and used unilateral *in vivo* channelrhodopsin-2 (ChR2) activation of VGluT2^+^ cells in the dmSC (Fig. 4a-c). In line with previous findings, activation of dmSC neurons in *Setd5^+/+^* animals instructed immediate shelter-directed escapes (Fig. 4d) and were absent in controls (Fig. S5). Gradually increasing the laser power and, thus, optogenetic activation, led to corresponding decreases in the latency to respond and an increase in the escape vigour in *Setd5^+/+^*(Fig. 4e). By contrast, activation of dmSC neurons in the *Setd5^+/-^* background more often resulted in an initial arrest, followed by shelter-seeking escape responses (Fig. 4f,g), mimicking the behavioural divergence observed for LER (Fig. 1). Interestingly, this behavioural difference was evident only during strong optogenetic activations. At lower laser intensities, both genotypes exhibited a transition from arrest to weak escape behaviours (Fig. 4h) with similarly low reaction time (Fig. 4e,g) and vigour (Fig. 4h). Correspondingly, the escape probabilities were identical at low optogenetic stimulation, but diverged with increased laser power (Fig. 4i). Given that changes in the strength of optogenetic activation mimic LER to looms of different contrast[29], we hypothesised that when probing animals to LER of different contrasts, both genotypes should show similar responses at lower contrast (Fig. 4j). To test this, we performed the LER at three different contrast levels: 98 %, 50 %, and 20 % (Fig. 4k,l). As predicted, the LER to low-contrast looms did not differ across genotypes, showing similar reaction times (Fig. 4k). Remarkably, in *Setd5^+/-^*animals, the reaction time remained largely constant across contrast levels (Fig. 4f,g), suggesting that the time required to initiate a response is independent of stimulus intensity. Together with the persistence of the arrest behaviour even at high levels of optogenetic activation (Fig. 4h), this supports the idea that there is a functional bottleneck downstream of the sensory processing circuits within the SC that impairs the timely generation of escape responses.

**Figure 4.**
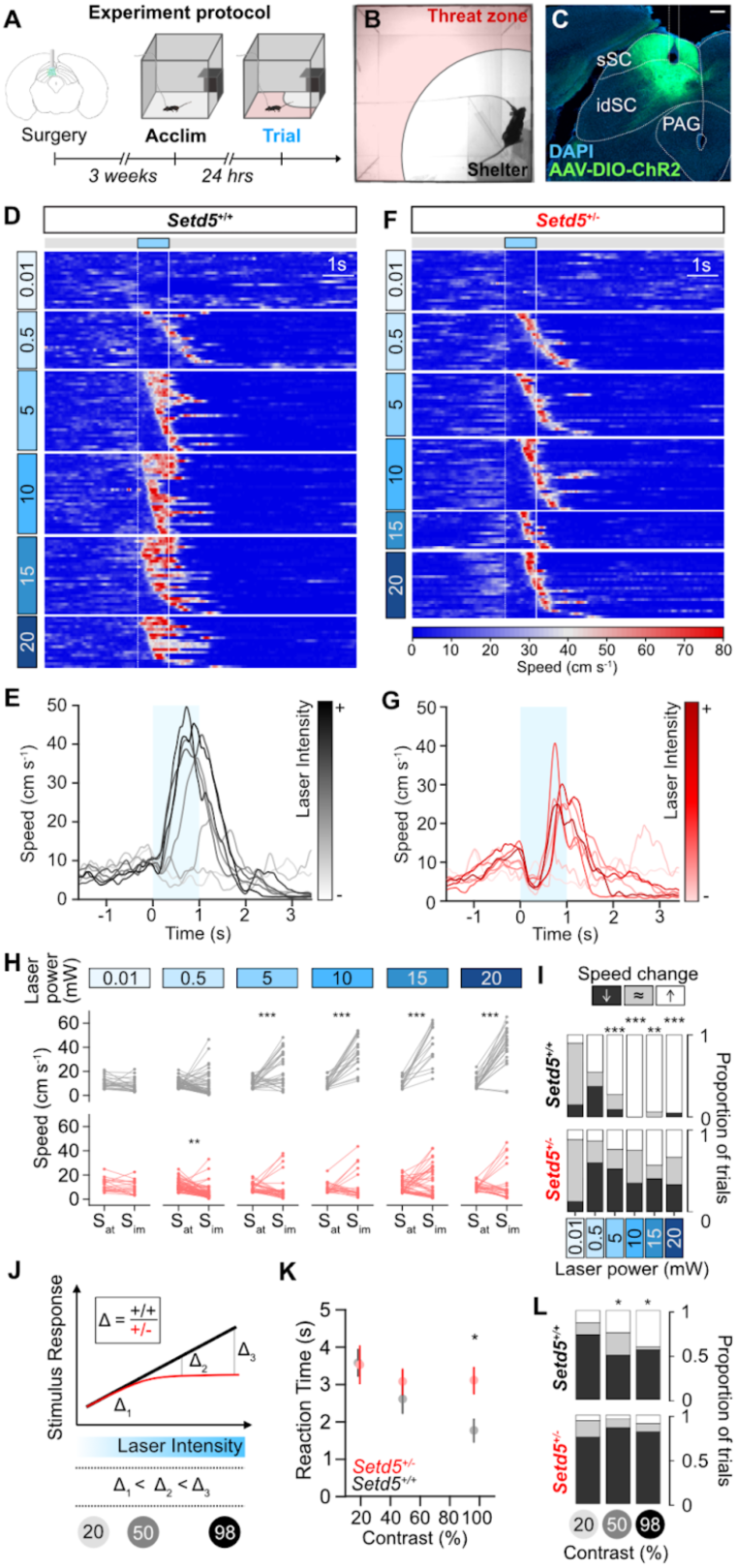
Activation of deep SC neurons recapitulates delayed LER in *Setd5^+/-^* animals. **a**, Timeline of the experimental protocol for optogenetic activation of the dmSC. **b**, Video frame during an optogenetics trial. **c**, Confocal micrograph of AAV-ChR2 expression in dmSC and optic fibre location reconstruction, scale bar: 200 μm. **d**, Raster plot of mouse speed in response to optogenetic activation, sorted by laser intensity and (**e**) mean speed responses at increasing laser intensities for *Setd5*^+/+^ (n = 4, 312 trials). **f-g**, as (**d-e**) but for *Setd5*^+/-^ mice (n = 4, 291 trials). Blue-shaded areas represent the laser stimulation. **h**. Subplots of trials in (**d, f**) showing the immediate change in speed upon light activation at different laser intensities for *Setd5*^+/+^ (n = 4, paired *t*-tests) and *Setd5*^+/-^ (n = 4, paired Wilcoxon’s tests) optogenetics trials. Sat is the mean speed of the animal ±50 ms of laser onset and Sim is the mean speed of the animal 300-800ms after laser onset. **i**, Proportion of trials at different laser intensities that either show an increase (white), decrease (black) or no change (grey) in speed upon light activation for *Setd5*^+/+^ (top) and *Setd5*^+/-^ (bottom). 0.01mW mm^-2^: 35 trials, p = 0.649; 0.5mW mm^-2^: 111 trials, p = 0.193; 5mW mm^-2^: 50 trials, p < 0.001; 10mW mm^-2^: 49 trials, p < 0.001; 15mW mm^-2^: 61 trials, p = 0.004; 20mW mm^-2^: 49 trials, p < 0.001, X^2^ test of independence. **j**, Schematic of the behavioural divergence between genotypes with increasing stimulus intensity (laser or loom). **k**, Summary of mean ± s.e.m. of reaction time to the LER paradigm at different stimulus contrasts (p = 0.021 for the interaction between genotype and contrast, *Setd5*^+/+^, n = 13, 68 trials; *Setd5*^+/-^, n = 13, 59 trials, repeated measures ANOVA. p = 0.018 for 98% contrast, with multiple comparisons and Bonferroni correction. *Setd5*^+/+^, 24 trials; *Setd5*^+/-^, 22 trials). **l**, Proportion of trials at different contrast looms that show an increase (white), decrease (black), or no change (grey) in speed upon light activation (20%: p = 0.698; 50%, p = 0.026; 98%, p = 0.021, X^2^ test of independence).

### dPAG neurons are hypoexcitable in the *Setd5* mouse model

Since our previous results point towards a disruption in the circuits downstream of the SC, we decided to investigate if the activity of the underlying dPAG is affected in the *Setd5^+/-^* model (Fig. 4). To test if any synaptic or intrinsic properties of dPAG neurons are impaired, we performed whole-cell patch-clamp recordings in slices (Fig. 5a). First, using the same animals used for optogenetic stimulation *in vivo*, we tested if the synaptic properties of dmSC inputs to the dPAG are changed. 10 Hz *ex vivo* optogenetic activation elicited monosynaptic excitatory postsynaptic current (EPSC) with an average current of −244±9.65 pA in *Setd5^+/+^* and -228±11.8 pA in *Setd5^+/-^* (Fig. 5a,b). At this frequency, repeated stimulations showed no facilitation in both genotypes (Fig. 5b), indicating that dmSC inputs to the dPAG are unaltered. Next, we explored spontaneous neurotransmission by recording sEPSCs (Fig. S6a-e), supporting the view that the synaptic properties remain similar at a circuit level. Finally, we probed the intrinsic properties of dPAG neurons. These experiments were done blind to the cell type, but we were able to classify them based on their firing statistics as excitatory and inhibitory[32] (Fig. S6f-j). Although the membrane potential, input resistance, membrane constant, and capacitance were identical between genotypes (Fig. 5c), we observed a stark decrease in the ability of these cells to generate action potentials in response to injected current in both excitatory and inhibitory neurons (Fig. 5d, Fig. S6h), without affecting the rheobase (Fig. S6k). These changes were accompanied by changes in the spike shape of Setd5*^+/-^* animals (Fig. 5e,f) that showed slightly higher depolarisation, lower after-hyperpolarisation and slower rise dynamics in comparison to their WT siblings (Fig. S6i,j). Based on prior work, we know that stronger dPAG activation leads to more vigorous escapes and that the underlying biophysical mechanisms depend on dPAG neuronal synaptic integration and excitability[29]. Thus, intrinsic excitability changes in excitatory neurons are a proxy of the expected escape strength. Remarkably, the response characteristics of dPAG neurons in *Setd5^+/+^ and Setd5^+/-^*match their behavioural differences (Fig. 1). At weak current-injections (Fig. 5d), optogenetic activation (Fig. 4d-h) or low contrast looms (Fig. 4k), both genotypes are indistinguishable from each other. However, at stronger current-injections *Setd5^+/+^ and Setd5^+/-^* diverge, with an hypoexcitability phenotype for *Setd5^+/-^*(Fig. 5d) that matches the delay in LER for optogenetic (Fig. 4d,f) and visual (Fig. 1, Fig. 4k) activations. To determine if the hypoexcitability phenotype was present in the preceding visual pathway, we patched dmSC interneurons (Fig. S6l-q). Here, no statistical significance was observed despite a hypoexcitability trend, indicating that the strength of the effect is area-specific but not ruling out other brain-wide deficiencies. In combination with previous work[29] and our optogenetic experiments (Fig. 4), these results suggest intrinsic deficits in the dPAG account, at least partly, for the delayed LER phenotype in the *Setd5*^+/-^ mice (Fig. 1).

**Fig. 5.**
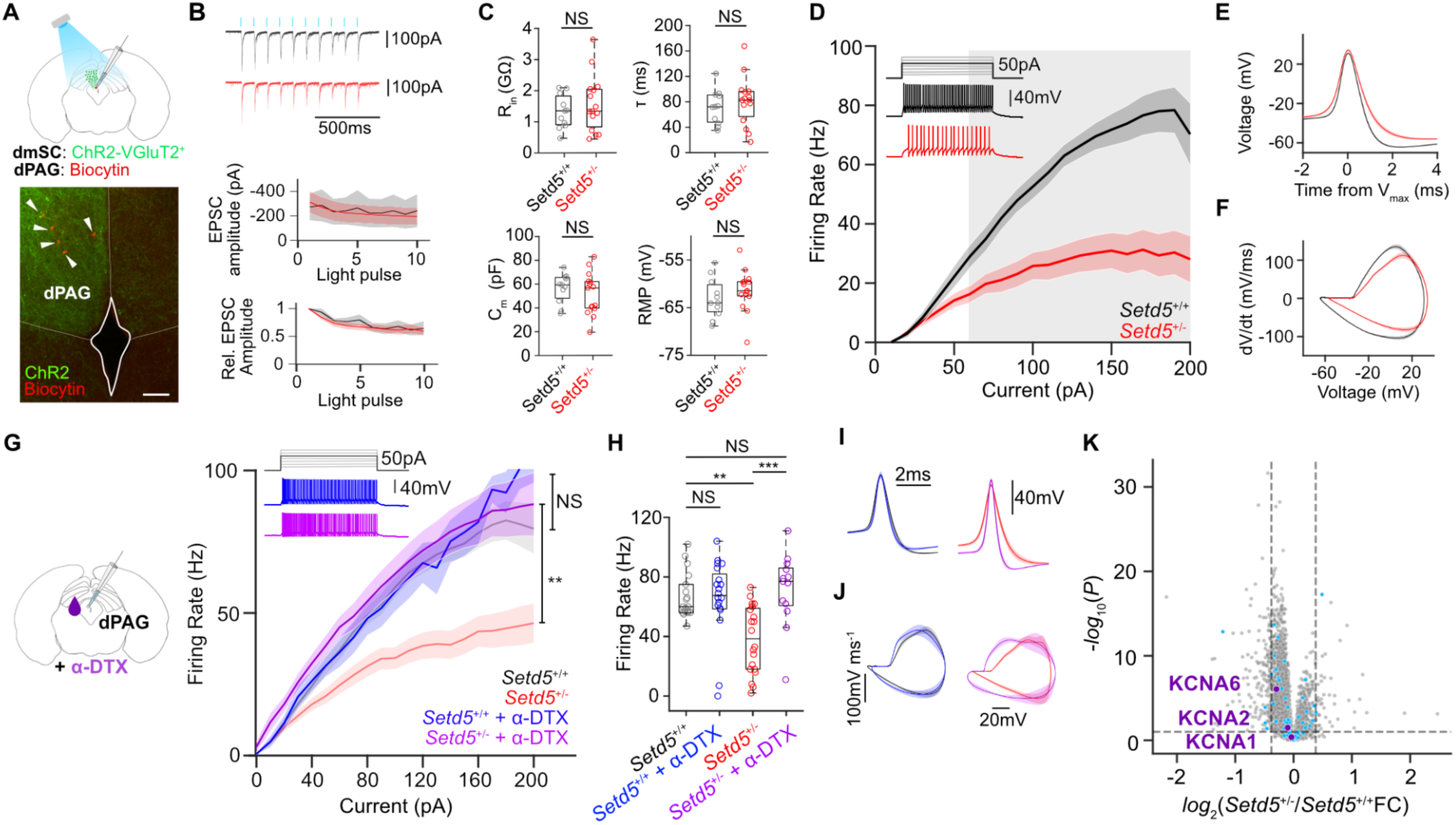
*Setd5*^+/-^ dPAG cells are hypoexcitable. **a**, Schematic of the experimental approach, *in vitro* patch clamp recordings (top) and micrograph of VGluT2^+^ dmSC projections to the dPAG infected with AAV9-ires-DIO-ChR2 (green) with dPAG biocytin filled and recorded cells (red & arrow heads, scale bar: 100 **µ**m). **b**, Top, whole-cell voltage clamp traces of example *Setd5*^+/+^ and *Setd5*^+/-^ cells (black and red, respectively) responding to 10 Hz light stimulation (blue ticks). Amplitude (middle, p = 0.078) and relative EPSC amplitude (bottom, p = 0.565) of responses to sequential light pulses in a 10 Hz train. **c**, Intrinsic properties of *Setd5^+/+^* (n = 6, 11 cells) and *Setd5*^+/-^ (n = 7, 14 cells) dPAG cells. Input resistance (P = 0.756, top left), membrane constant tau (P = 0.436, top right), membrane capacitance (P > 0.995, bottom left) and resting membrane potential (P = 0.213, bottom right). **d**, Summary of the relationship between current injection and action potential firing for putative glutamatergic cells showing a strong reduction in firing (p < 0.001). Inset, representative example traces to a 50 pA current injection. Grey area indicates the current injection values that significantly differ between *Setd5*^+/+^ cells (black) and *Setd5*^+/-^ cells (red) found by a multiple comparisons analysis with Tukey correction. **e**, Average shape and (**f**) phase plane analysis of the action potentials generated in the rheobase sweep (*Setd5*^+/+^, 13 cells, 42 spikes; *Setd5*^+/-^, 14 cells, 72 spikes). **g**, Summary of the relationship between current injection and action potential firing for all *Setd5*^+/+^ cells (black, n = 18) and *Setd5*^+/-^ cells (red, n = 20) before and after (*Setd5*^+/+^: blue, n =12 cells; *Setd5*^+/-^: purple, n = 13 cells) application of **α**-Dendrotoxin (**α**-DTX, 100 nM, p > 0.995 and p = 0.0147 for the effect of **α**-DTX on *Setd5*^+/+^ and *Setd5*^+/-^ firing, respectively). Inset, representative example traces from *Setd5*^+/+^ cells (blue) and *Setd5*^+/-^ (purple) cells after **α**-DTX application to a 50 pA current injection. **h**, Effect of **α**-DTX on firing in response to 120pA current injection. Before **α**-DTX (*Setd5*^+/+^ versus *Setd5*^+/-^; p = 0.0017), effect of **α**-DTX on *Setd5*^+/+^ (before versus after **α**-DTX; P > 0.995) and *Setd5*^+/-^ (before versus after **α**-DTX; P < 0.001, *Setd5*^+/+^ before versus *Setd5*^+/-^ after **α**-DTX; P > 0.995). Multiple comparison analysis after rm-ANOVA with Tukey correction. **i**, Action potential shape and **j**, phase plane analysis of the action potentials generated in the rheobase current in *Setd5*^+/+^ cells (without **α**-DTX, black, n =19; with **α**-DTX, blue, n =12), *Setd5*^+/-^ cells (without **α**-DTX, red, n=21; with **α**DTX, purple, n = 13). **k**, Volcano plot for differential protein levels between adult S*etd5*^+/+^ and *Setd5*^+/-^ mice (n = 6 samples per genotype, cyan dots represent proteins annotated as ion-channels and purple dots represent Kv1.1, Kv1.2 and Kv1.6. Lines and shaded areas, mean ± s.e.m., respectively. Box-and-whisker plots show the median, IQR, and range. P-values are Wilcoxon’s rank sum test. p-values are two-way repeated measures ANOVA. Volcano plot horizontal dashed line represents the significance threshold (P-value < 0.1) from a two-sided moderated t-test while the vertical dashed lines indicate fold change values greater or lower than 0.4 between *Setd5*^+/+^ and *Setd5*^+/-^.

#### dPAG cells are hypoexcitable due to K_v_ channel misregulation

Next, we investigated the underlying mechanisms of the hypoexcitable phenotype of the *Setd5^+/-^* dPAG cells. The changes in excitability and spike kinetics are indicative of differences in channel composition that are part of adaptive homeostatic mechanisms[33], suggesting the influence of voltage-gated potassium channels (Kv). Previously published work[23], which studied the transcriptomic changes brought about by a mutation in *Setd5^+/-^,* highlighted significant upregulation in the expression of Kv1.1 channels during development. To test whether the blockade of Kv1.1 would rescue the hypoexcitability phenotype, we applied 100 nM of **α**-Dendrotoxin (**α**-DTX), a specific Kv1.1, Kv1.2 and Kv1.6 blocker, and patched dPAG cells. While **α**-DTX had no effect on *Setd5^+/+^*excitability of dPAG cells, strikingly, *Setd5^+/-^* cells reversed their hypoexcitability phenotype to WT levels (Fig. 5g-h). In addition, after **α**-DTX application, the spike dynamics remained identical in *Setd5^+/+^*, whereas *Setd5^+/-^*became faster, mimicking the WT properties (Fig. 5i-j). These differences are not due to an overexpression of Kv channels. Tissue-specific proteomic analysis showed no difference in the protein level of Kv channels in either the PAG (Fig. 5k), SC (Fig. S7a) or cortex (Fig. S7b). This is in accordance with immunohistochemical (Fig. S7c-f) and western-blot analysis (Fig. S7g,h). Given that Kv1.1 has been recently shown to be involved in the homeostatic control of firing[33,34] through modulation of the axon initial segment (AIS)[35], these results indicate that the deficits might not arise due to direct changes in expression but due to homeostatic misregulation. Next, we tested if the dPAG neurons of *Ptchd1^Y/-^* and *Cul3^+/-^* animals showed similar physiological properties to *Setd5*^+/-^ neurons. As expected from the molecular differences among the tested ASD models, the physiological characteristics differed. *Ptchd1* animals, despite having mostly identical intrinsic properties to their WT siblings (Fig. S8a-d), also show a hypoexcitability phenotype at high current inputs (Fig. S8e-i,s). However, this hypoexcitability phenotype cannot be rescued by **α**-DTX application (Fig. S8s). *Cul3* animals did not show any changes in either their current-firing relationship or intrinsic properties (Fig. S8j-r,u,v). Overall, these results indicate that different models might have distinct neurodevelopmental origins or key molecular changes that give rise to particular behavioural traits. In particular, for *Setd5,* Kv1 channels appear as key players and targets to reverse the delayed LER phenotype and probe the causality of its correlation with place avoidance deficits.

To investigate whether the area-specific dPAG hypoexcitability phenotype mediated the delayed LER, we tested whether **α**-DTX application targeted directly to the dPAG could rescue the behavioural phenotype (Fig. 6a). We implanted cannulas above the dPAG in *Setd5^+/-^*and their WT littermates, a target-specific approach as visualised through neurobiotin injections (Fig. 6b). Initially, we injected saline as a control in both cohorts and subsequently tested the LER (Fig. 6c,d, top). As previously demonstrated (Fig. 1), *Setd5^+/-^*animals responded slower than their WT littermates and had no difference in their maximal escape speed during the first presentation day (Fig. 6d, top). However, the behaviour of both cohorts were indistinguishable from each other when **α**-DTX was applied, escaping immediately after the loom presentation (Fig. 6c,d, bottom, Video S3), with similar reaction times (Fig. 6e) and vigour (Fig. 6f), even despite *Setd5^+/-^* animals normally adapting across days (Fig. 2d). We then quantified the shelter exits of both cohorts (Fig. 6g,h) to investigate whether the correlation between shelter exits and delayed LER (Fig. 2e) could have a common origin. Remarkably, *Setd5^+/-^* animals recovered their place avoidance as well after **α**-DTX application (Fig. 6h), indicating that the timely initiation of actions and memory formation are closely linked. These results suggest that perceptual deficits can have far-reaching consequences for unrelated behavioural traits.

**Fig. 6.**
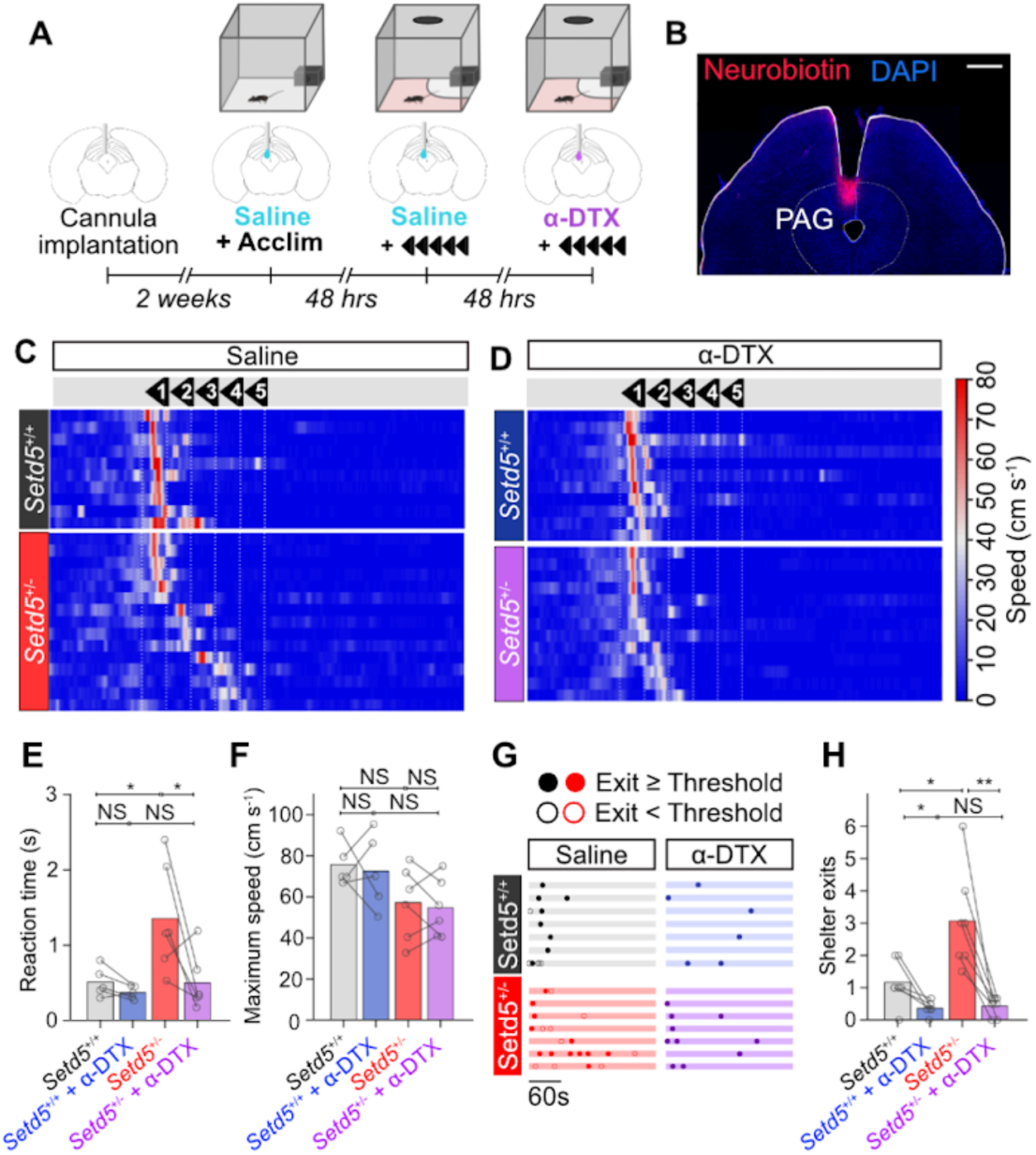
Pharmacological Kv channel block rescues delayed LER and place avoidance. **a**, Timeline of *in vivo* **α**-DTX cannula experiments. **b**, Confocal micrograph of a coronal section of the location of the cannula above the dPAG as well as the expression of neurobiotin that was infused into the cannula at the end of the experiments. Scale bar: 500 **µ**m. **c**, Raster plots of mouse speed in response to the looming stimuli for *Setd5^+/+^* before (top, n = 6, 10 trials) and after (bottom, n = 6, 11 trials) infusion of **α**-DTX (500nL, 500nM) sorted by reaction time. **d**, as for (**c**) but for *Setd5*^+/-^ before (top, n = 6, 15 trials) and after (bottom, n = 6, 13 trials) animals after infusion of **α**-DTX (500nL, 500nM). **e**, Effect of **α**-DTX on reaction time. Before **α**-DTX (*Setd5*^+/+^ versus *Setd5*^+/-^; P = 0.017), effect of **α**-DTX on *Setd5*^+/+^ (saline versus **α**-DTX; p = 0.104) and *Setd5*^+/-^ (saline versus **α**-DTX; p = 0.014) and after **α**-DTX (*Setd5*^+/+^ versus *Setd5*^+/-^; P = 0.571). **f**, Effect of **α**-DTX on escape vigour. Before **α**-DTX (*Setd5*^+/+^ versus *Setd5*^+/-^; P = 0.052), effect of **α**-DTX on *Setd5*^+/+^ (saline versus **α**-DTX; p = 0.760) and *Setd5*^+/-^ (saline versus **α**-DTX; p = 0.649) and after **α**-DTX (*Setd5*^+/+^ versus *Setd5*^+/-^; P = 0.075). **g**, Shelter exit behaviour during the first LER trial when the mice are injected with saline (*Setd5*^+/+^, black, top left; *Setd5*^+/-^, red, bottom left) or **α**-DTX (*Setd5*^+/+^, blue, top right; *Setd5*^+/-^, purple, bottom right). Each row represents one animal, filled and open dots represent exits crossed into the threat zone or not, respectively. **h**, Effect of **α**-DTX on shelter exits. Before **α**-DTX (*Setd5*^+/+^ versus *Setd5*^+/-^; P = 0.012), effect of **α**-DTX on *Setd5*^+/+^ (saline versus **α**-DTX; p = 0.038) and *Setd5*^+/-^ (saline versus **α**-DTX; p = 0.003) and after **α**-DTX (*Setd5*^+/+^ versus *Setd5*^+/^; P = 0.495). Markers represent the average values across all trials for individual animals. P-values: Wilcoxon’s test, p-values: paired-t-test.

## Discussion

Autism spectrum disorders (ASDs) are widespread in the global population, but our understanding of the disorder’s origins remains limited. Advancements in genetic sequencing technologies have enabled the isolation of autism-risk genes, opening the door to several in-depth studies into the molecular mechanism of action of these genes[36]. However, there is a considerable mismatch between the observed wealth of molecular changes and our understanding of their roles in changing circuit function and, correspondingly, behaviour. The latter is particularly relevant since ASD is diagnosed by a combination of behavioural traits known as diagnostic criteria[2]. Although several behavioural paradigms are being studied across mouse models[37], the variability and experimental intricacies of behavioural studies make direct comparisons across models difficult. This has led to studies focusing on single models, defining particular changes but not general principles[38]. Furthermore, these studies have primarily centred on the intricacies of social interactions, communication, and repetitive behaviours—complex traits that present a formidable challenge in establishing a clear connection between neuronal dysfunction and behavioural manifestations. Here we show that the study of innate defensive behaviours, a perceptual task, provides a convergent behavioural framework that enables the directed analysis of the underlying sensorimotor processes up to the molecular level, permitting comparative dissections of the neuronal dysfunctions across ASD models and a systematic understanding of the neuronal mechanisms underlying the co-occurrence of behavioural traits. This is particularly interesting as the severity of sensory and perceptual impairments has been strongly linked to the strength and expression of traditional, non-sensory ASD traits [18–20].

Defensive escape behaviours are among the most fundamental perceptual decisions performed by animals [22]. Their finely-tuned mechanisms are indispensable for survival, but also for properly interacting with the environment. Whereas some stimuli unambiguously signal an imminent threat and should instruct immediate action, others are ambiguous and require adaptive responses arbitrated by the current context and state to decide, e.g., between ignoring, freezing, fighting, or escaping[22]. In mice, behavioural escape decisions are known to be initiated by the dPAG[29], where appropriate escape decisions are thought to be determined. While all tested ASD models can, in principle, respond behaviourally as robustly as their WT siblings, they require longer and respond with less vigour once an action has been initiated, hampering their ability to develop an appropriate place avoidance to the threat zone (Figs. 1-3). The underlying changes were rigorously dissected in the *Setd5* haploinsufficient model, pointing to a specific misregulation of voltage-gated potassium channels in dPAG neurons that gives rise to a strong hypoexcitability phenotype (Figs. 3-5), namely Kv1.1, Kv1.2 and Kv1.6. Targeted pharmacological rescue of this hypoexcitability *in vivo* completely reverses the behavioural phenotype (Fig. 6). This is an important finding since other brain areas are also known to be required for proper defensive responses, in particular, freezing, such as the pathway to the basolateral amygdala (BLA) via the lateral posterior thalamus (LP)[9,39]. Our results indicate a specific involvement of the dPAG and not the BLA via LP pathway.

In addition to the delayed LER, we observed a strong reduction in place avoidance to the threat zone that elicited maladapted repetitive behaviour (Fig. 2, Suppl. Video 2). This suggests that ASD mouse models either have deficits in fear memory formation or are intrinsically less fearful, thus, incapable of appropriately interpreting the noxiousness of the threatening experience (Figs. 1, 2, Video S1, S2)—maladaptive behaviours that recapitulate some human ASD traits[40]. Interestingly, this reduction of place avoidance is directly linked with longer reaction times to the LER across ASD models (Fig. 2e, 3h, p), and thus, with the dPAG hypoexcitability phenotype (Fig. 5d). We show that this correlation is causally linked, given that the target-specific pharmacological rescue reverses both, the delayed LER and place avoidance to wild-type levels (Fig. 6, Suppl. Video 3). Thus, maladaptive perceptual decisions, as with the delayed LER, can profoundly affect seemingly independent behavioural traits, such as LER, threat-induced place avoidance and the emergence of repetitive behaviours (Videos S1 & S2). This is in line with studies in humans. Threat imminence has been shown to elicit PAG activation[41] and the electrical stimulation of midbrain structures elicited strong emotional reactions[42]. In rodents, the dPAG has been shown to support fear learning[43], particularly in contextual conditioning paradigms[44]. Overall, our results emphasise the role of subcortical pathways through the PAG in the altered perceptual abilities frequently described in ASD, and fear memory formation in general.

The dPAG hypoexcitability phenotype indicates that the integration and action initiation required for adequate perceptual decision-making is disrupted in *Setd5* animals. This physiological phenotype is tightly linked with behavioural impairments. Substantial threat evidence, either by strong optogenetic activation of dmSC neurons or high-contrast visual looms, elicit a slower and less robust response in *Setd5^+/-^*compared to their WT siblings. On the other hand, limited threat evidence, either by weak optogenetic activation or low-contrast looms, instructs similar behavioural responses between genotypes (Fig. 4). Accordingly, only strong current-injections that match the expected dPAG drive caused by high contrast loom[29] cause a Kv-channel induced hypoexcitability phenotype (Fig. 5), indicating that the delayed response phenotype is due to a dPAG dysfunction. This stimulus strength relationship aligns with the coping difficulties of many individuals with ASD to salient sensory stimuli, such as bright lights or crowded places, but not to mellow sensory environments[6]. Recently, changes in dPAG excitability and the expression of defensive behaviours have also been identified in the *Nlgn3*^+/-^ rat model of autism [42]. Notably, these rats exhibit the inverse behavioural and physiological phenotype to that observed in *Setd5*^+/-^ mice, displaying stronger responses auditory fear conditioning, prolonged place avoidance and hyperexcitability in dPAG cells. These complementary results reflect the wide range of sensory sensitivities observed in the human population with ASD, and reinforce the PAG as an important area to investigate disrupted sensory processing and its far-reaching effects in the core symptomatology associated with autism.

Given that the expression levels of Kv channels remain unaltered (Fig. 5k, Fig. S7), the causes of the physiological hypoexcitability phenotype probably arise through homeostatic misregulation, either by localisation[33], post-translational modifications[45], or interactions with auxiliary subunits[46]. The link between potassium channels and autism is well established[47]. Genetic analyses of individuals with ASD uncovered deleterious mutations in potassium channels[47]. These, however, exclude Kv1.1. However, autistic-like repetitive and social behaviours in the *Scn2a* haploinsufficiency mouse model have been rescued in a Kv1.1 deficient background[48], supporting our results that enhancing and not disrupting Kv1.1’s contribution can be a fundamental factor in ASD.

Sensory processing and perceptual abnormalities have been shown to co-occur with classical ASD diagnostic criteria and have been recently proposed as promising behavioural biomarkers of autism[4]. Here we show that the innate looming escape response (LER) provides a reliable and quantitative framework to systematically link behavioural traits with the underlying molecular changes. The exact molecular alterations are dependent on the model. Whereas *Setd5* and *Ptchd1* dPAG neurons show a hypoexcitability phenotype via different mechanisms, the weaker LER deficits in *Cul3* are independent of dPAG intrinsic properties (Fig. S8). Importantly, we show that in the *Setd5* haploinsufficient model, these behavioural traits are not necessarily developmental since they can be ameliorated pharmacologically in adulthood. This is an encouraging result that may reveal therapeutic targets as envisioned by precision medicine approaches[36].

In summary, our work links innate LER dysfunctions across diverse genetic mouse models of ASDs. Specifically, in the *Setd5* haploinsufficient model, we found that a key behavioural node, the PAG[49], functions as an interface between sensory, limbic and motor circuits that, when disrupted, causally affects seemingly unrelated behaviours. This appears to be related to observations in people with ASD, where the severity of sensory processing impairments is related to the severity of core symptoms associated with autism[18–20]. Future studies designed to dissect the causal relationships between innate sensorimotor deficits, such as LER, and core ASD behavioural symptoms, such as social and communication difficulties, will be a revealing avenue to build a comprehensive view of ASD.

## Materials and Methods

### Mice

The study was discussed and approved by the institutional ethics committee of the University of Veterinary Medicine Vienna and the Institute of Science and Technology Austria in accordance with good scientific practice guidelines and national legislation under license numbers BMWF-68.205/0023-II/3b/2014 and 66.018/0017-WF/V/3b/2017. Male and female adult *Setd5*^+/-^, *Setd5*^+/fl^::*Emx1*-Cre, *Setd5*^+/-^::*VGluT2*-ires-Cre, *Cul3*^+/-^, *Ptchd1*^Y/-^ mice were housed under a 12h light/dark cycle (lights on at 07:00), with food and water available ad libitum. The animals were housed in groups of 2-6 animals per cage and were tested during the light phase. Animals used for the *in vivo* electrophysiological recordings or freely-moving optogenetics experiments were group housed before surgery and individually housed after the surgery. Mice were chosen based on genotypes. Sex-matched animal pairs of control-mutant siblings from the same litters were compared to decrease variance due to age, environment and genetic background.

### Generation of Ptchd1^Y/-^ mice

*Ptchd1*^Y/-^ mice for analysis were generated using the CRISPR/Cas9 system, as described in[50]. In brief, *in vitro* transcribed sgRNAs targeting the exon 2 of *Ptchd1* were microinjected into zygotes of C57BL/6NRj mice together with Cas9 *mRNA*. Offspring were screened by PCR for deletions using primers spanning the target sites (5′-GTAGGGCTGGAATCATGAGG-3′, 5′-CACATCCTTTGGTGTGATGC-3′). Deletion of *Ptchd1* exon 2 was confirmed by Sanger sequencing analyses. The founder line was established from a female *Ptchd1*^+/-^ mouse with a 1.26 kb deletion. Analysis of *Ptchd1* transcript level by RT-PCR confirmed impaired *Ptchd1* expression in brain, cerebellum and kidney of the *Ptchd1*^Y/-^ mice, compared to control mice.

### Behavioural procedures

#### Experimental setup

All behavioural experiments were performed at mesopic light levels, in a square, black, IR-transparent Perspex box (W: 32 cm x L: 32 cm x H: 32 cm) with a shelter made of the same material (9 cm x 9 cm x 3 cm) positioned in one corner. One wall of the box was 2 cm shorter to allow for ventilation and for the optic fibre to pass through for optogenetic experiments. A plastic screen covered with a plastic sheet with a matte surface was placed as a lid on top of the box, and a modified Texas Instruments DLP projector (DLP CR4500EVM) back-projected a grey background. The blue LED was exchanged for a high-power UV-LED (ProLight 1W UV LED, peak 405 nm) to improve the differential stimulation of S pigments. The experiments were recorded with a near-IR camera (acA1920-150um, Basler, 60 Hz) from below the centre of the arena. Video recording, visual and optogenetic stimulation were controlled using custom software written in MATLAB, python and Arduino. The entire apparatus was housed within a sound-deadening, light-proof cabinet which was illuminated by an infrared light-emitting diode (LED) lamp (SIL 616.21, Sanitas).

#### Standard looming escape response (LER) protocol

All animals were habituated to the test arena at least one day before testing and were allowed to explore the arena for at least 20 mins. The initial 10 min of this exploration time was recorded and used to analyse the animals’ exploratory behaviour (Fig. S1). For the looming avoidance response (LER: Figs.1, 2, 3, 6, S1, S2, S3) paradigm, responses were tested over five consecutive days. On each test day the animals were given 10 min to acclimatise to the arena, after which they were subjected to 3 test trials, with a 3 min interval between each trial. Each trial lasted 180 s, within which the central position of the mouse was tracked online (Python) and was used to trigger the visual stimuli in a closed-loop manner whenever the mouse crossed a designated threshold distance (10 cm) from the centre of the shelter, with a minimum stimulus interval of 30 s. A typical experiment lasted ∼30 min and the entire arena was cleaned with 70 % ethanol between animals.

#### LER protocol with a food reward

In preparation for the reward trials, after the 4^th^ day of testing a small piece of banana chip was placed in the opposite corner of the arena to the shelter and the mice were free to explore the arena without triggering any loom stimuli. The animals were left to explore until they successfully acquired the banana chip, after which time both the mouse and the chip were placed back in their home cage. The reward trial then took place after the 5^th^ day of testing. Another small piece of banana chip was again placed in the opposite corner of the arena and the animals were free to explore, however this time whenever the mouse passed the threshold distance from the shelter a looming stimulus would start, this time with no inter-stimulus interval. For different contrast experiments, animals were presented with looming stimuli of 20, 50 or 98 % contrast in a pseudorandom order upon crossing the trigger threshold. The background luminosity remained constant while the luminance of the disc was increased to generate low contrast stimuli.

#### LER protocol with different contrast stimuli

Mice were similarly habituated to the arena as in the standard LER experiments; however, animals underwent a single session of three LER tests over one day. In contrast to the standard LER protocol, in these experiments, upon entering the threat zone they were exposed to looming stimuli of 20, 50 or 98% contrast in a pseudorandom order. The background luminosity remained constant while the luminance of the disc was increased to generate low contrast stimuli.

#### In vivo optogenetic activation experiments

The same behavioural arena was used for the optogenetic activation experiments except with the standard shelter exchanged for an open shelter consisting of a black piece of perspex (9 cm x 5 cm) positioned 10 cm above the floor. Mice were similarly acclimatised to the arena, as in the LER procedure, but with the fibre optic cable (200 μm, 0.48 NA, Doric Lenses) attached via a rotary joint (RJ1, ThorLabs) to allow for unrestrained movement and minimal handling of the animals. Since it has previously been shown[29] that increasing light intensity can be used as a proxy for the level of mSC, and hence dPAG, activation we decided to systematically modulate the laser light intensity in different trials. Once the trial began, mice were photostimulated (473 nm, 10 light pulses of 10 ms at 10 Hz; Shanghai Dream Lasers Technology) if they passed the threshold distance (10 cm) from the shelter, with a minimum inter-stimulus interval of 30 s. The initial laser intensity was set to a low irradiance (0.01-0.1 mW mm^-2^) that did not evoke an observable behavioural response, then increased to 0.5, 5, 10, 15 and 20 mW mm^-2^ with at least 3 tests (180 s each, with a maximum of 5 trials per test) at each intensity. Two mice never elicited an observable escape behaviour, and post-processing histological analysis revealed mislocalised or the absence of viral expression in these animals and they, along with their matched siblings, were excluded from the analysis.

#### In vivo cannula experiments

The same behavioural arena was used for the cannula experiments with ɑ-dendrotoxin (ɑ-DTX). Two weeks after mice were implanted with the cannula, they were acclimatised to the arena for 20 minutes and gently handled. 48 hours after acclimatisation to the arena, the mice were attached to the infusion system (tubing (PlasticsOne C313CT), infusion cannula (C315IS-5/SPC, 33GA, 5mm), guide cannula (PlasticsOne, C315GS-5/SF, 26GA, 5mm) and a 0.5 μL manual syringe (Model 7000.5, #86250, Hamilton) and placed into an open cage to allow them to move freely during the infusion of 500nL of saline with 0.1 % bovine serum albumin (BSA) at 100nl min^-1^. The mice were left for a further 5 min before detaching them from the infusion system and placing them within the arena. The first 5 minutes the mice were within the arena was recorded to assess the baseline activity of these animals. 15 minutes after the start of infusion, a single trial of the standard LER trial was initiated. 48 hours after the first LER trial with saline, the mice were reattached to the infusion system and ɑ-DTX (500nL, 500nM in 0.1 % BSA) was infused at 100nl min^-1^. The mice were again left for 5 minutes before being detached and placed into the arena. Baseline activity was assessed by recording the first 5 minutes within the arena, then 15 minutes after the start of the infusion the standard LER trial was initiated. This time, there was a 3 min gap between LER trials and 3 trials were conducted, as in the standard LER test. After the last test, the mice were re-attached to the infusion system and infused with 500nL of a 10% solution of neurobiotin in sterile saline at 100nl min^-1^, left for 5 minutes, then detached and placed back into their home cages. The animals were sacrificed 48 hours after infusion with neurobiotin and later stained with Streptavidin-594 in order to confirm the location of the cannula implantation and infusion.

### Viruses

The viruses used in this study: for optogenetic activation, adeno-associated virus (AAV) AAV9-EF1a-DIO-hChR2(E123T)T159C)-EYFP-WPRE-hGH (1×10^¹³^ viral genomes per ml (vg ml^-1^), Addgene 35509); for control experiments, AAV9-Syn-Flex-GCaMP6m-WPRE-SV40 (1×10^¹³^ vg/ml, Addgene 100838).

### Surgical procedures

Mice were anesthetised with an intraperitoneal (i.p.) injection of ketamine (95 mg kg^-1^) and xylazine (4.5 mg kg^-1^), followed by metamizol (20 mg kg^-1^, i.p.), buprenorphine (0.05 mg kg^-1^, i.p.) and meloxicam (2 mg kg^-1^, subcutaneous (s.c.)). If needed, isoflurane (0.5-2 % in oxygen, 0.8 l min^-1^) was used to maintain anesthesia. Fur on the top of the head was removed with an electric hair remover (Braun Precision Trimmer, PT 5010) then the mice were placed in a stereotaxic frame (Model 962, Kopf Instruments) and fixed using the ear bars. Eyes were protected using Oleo Vital eye cream and a topical analgesic (Xylocain 2 % gel) was applied with a q-tip before cutting the skin. Craniotomies of ∼ 1mm diameter were made using 0.5 mm burrs (Fine Science Tools) and dental drill (Foredom, HP4-917) and viral vectors were delivered using pulled pipettes (World Precision Instruments, 1.4 mm OD, 0.53 mm ID, #504949) made with a micropipette puller (DMZ Zeitz-puller, Germany) and delivered using an automatic nanoinjector (World Precision Instruments, Nanoliter 2010) at 20 nl min^-1^. The skin was closed after surgery using surgical glue (Vetbond, 3M, #1469SB).

#### Optogenetics experiments

Setd5^+/+^::VGluT2-ires-Cre and Setd5^+/-^::VGluT2-ires-Cre sex-matched sibling mice were injected with AAV9-DIO-ChR2-eYFP (see Viruses) into the left dmSC (75-100 nl, ML: -0.5, AP: -0.5 to -0.7, DV: -1.45 to -1.6, from bregma). Control mice were injected with 100nl of AAV9-Flex-GCaMP6m into the left dmSC at the same coordinates. One optic fibre (400 μm diameter, CFMC54L02, ThorLabs) was implanted 250 μm dorsal to the injection site. Fibres were affixed using light-curing glue (Optibond Universal, Kerr Dental) and dental cement (SuperBond C&B Kit, Hentschel-Dental).

#### In vivo electrophysiological recordings

Mice were implanted with a custom-designed head plate over a ∼0.5-1 mm diameter craniotomy (left hemisphere, AP: 0.4 to -0.4, ML: 0.4 to 0.8 from lambda) and a second craniotomy (∼0.5 mm) was made anterior to bregma on the right hemisphere, where a reference gold pin was implanted and lowered until it touched the brain surface. Both were cemented to the skull using light-curing glue (Optibond Universal, Kerr Dental) and dental cement (SuperBond C&B Kit, Hentschel-Dental), with the headplate additionally fixed using Charisma Flow (Kulzer GmBH). The craniotomy was covered with Kwik-Cast (World Precision Instruments) to protect the underlying tissue before recording.

#### In vivo cannula experiments

Setd5^+/+^ and Setd5^+/-^ sex-matched sibling mice were implanted with a single infusion cannula (C315IS-5/SPC, 33GA, 5mm) above the dorsal PAG (ML: - 0.5, AP: -0.8 to -1, DV: -1.5 to -1.65, from lambda). Fibres were affixed using light-curing glue (Optibond Universal, Kerr Dental) and dental cement (SuperBond C&B Kit, Hentschel-Dental) and the skin was closed around the implant with surgical glue (Vetbond, 3M, #1469SB).

### Visual Stimuli

All stimuli were created and presented using custom-made scripts in MATLAB (MathWorks, Inc.), using the Psychophysics Toolbox extensions[51] and frames were presented at a refresh rate of 60 Hz.

#### Arena behavioural experiments

The standard stimulus consisted of a dark disc on a grey background presented on the lid over the centre of the arena, that reached a maximum size of 40 deg/visual angle. The disc expanded over 500 ms at a speed of 80 deg s^-1^ and then remained at this size for 250 ms before disappearing and a new expanding disc appeared immediately. A maximum of 5 expanding discs were shown upon triggering the visual stimulus. A red rectangle was presented along one side of the projected image every 5^th^ frame, onto an out-of-view section of the arena lid. An optical filter reflected this red light back towards a photodiode (PDA36A2, ThorLabs) and this information was then used for post-hoc synchronisation of the behavioural data and the stimulus.

#### Head-fixed in vivo electrophysiological experiments

Sensory stimuli were made using the Psychophysics Toolbox extensions in MATLAB (MathWorks, Inc.) and presented and synchronised using custom-made LabVIEW software. At each depth, the mice were exposed to one 20 min presentation of a full field shifting spatiotemporal white-noise stimulus (see [52]) in order to assess receptive fields, and two repetitions of the remaining stimuli (5 x loom bouts, moving bars, moving gratings, full-field flash). Each presentation would start with a uniform grey screen (60 s, green and UV LED intensities were set to match the sun spectrum from a mouse opsin perspective), followed by the first presentation of the looming stimulus, then the other stimuli in a pseudorandom order separated by 30 s gaps of a grey screen. White-noise “checker” stimuli were presented at 20 Hz update for 10 min. The checker size was 6.56 deg^2^ and the entire grid was shifted between 0 and 3.25 deg in both x-and y-axis after every frame. Auditory monitoring of responses to a flashing point stimulus was used in real-time to gauge the rough centre of the receptive fields of the neurons being recorded and the location of the looming stimulus was positioned to overlap with this position. The looming stimulus consisted of 10 repetitions of 5 sequential, individual looming stimuli; the same stimulus previously described and used in the behavioural experiments. These loom bouts would be presented with intervals of pseudorandom lengths between 10-30 s with a grey screen in between. To assess direction selectivity, the mice were presented with full-filed moving square gratings (1.5 cycles s^-1^ temporal frequency and 0.3 cycles deg^-1^ spatial frequency) or a wide dark bar (38 deg s^-1^ speed, 6.5 deg width) in four orientations, drifting in eight different directions (separated by 45 deg) repeated twice for each orientation. Full-field ON-OFF flashes were presented in blocks of 10 repetitions of 0.5 s OFF, 1 s ON and 0.5 s OFF stimuli.

### *In vivo* electrophysiology

#### Data acquisition

A 32-channel silicon probe (NeuroNexus, USA, A32-OM32) was used to record extracellular spikes from different depths of the SC in *Setd5*^+/+^ (n = 3), *Setd5*^+/-^ (n = 3), *Cul3*^+/+^ (n = 3), *Cul3*^+/-^ (n = 3), *Ptchd1*^Y/+^ (n = 3) and *Ptchd1*^Y/-^ (n = 3) sex-matched sibling mice. At least 24 hrs after surgery, mice were placed on a suspended spherical treadmill (20 cm diameter) and head-fixed within a toroidal screen covering 190 ° of visual angle which was illuminated by a DLP-light projector (Blue LED exchanged for UV 405 nm, 60 Hz) reflecting off a spherical mirror in front of the screen (see diagram in Fig. S4a). The behaviour of the animal was recorded by a near-IR camera (Basler, acA1920-150um, 30 Hz) from the reflection of the animal in an IR-reflective heat mirror positioned 30 cm from the face of the mouse and 35 cm from the camera. Prior to recording, the probe was coated with DiI (1 mM in ethanol, Merck, #42364) for later histological confirmation of the recording site and a wire was connected to the ground pin for both an external reference and ground. The kwik-cast seal was removed, revealing the brain surface and the confluence of the overlying superior sagittal and the left transverse sinus. Care was taken to position the probe posteriorly and medially as possible without damaging the sinuses or other blood vessels. It was then zeroed at the brain surface then slowly lowered into the overlying cortex and the SC at 1 mm s^-1^ to depths between 1.2-2.2 mm. The recording was started 15 min after reaching the desired depth to allow for stabilisation. The probe was lowered to 3-4 depths per animal with 250 μm intervals, to sample from cells across the depth of the SC. Each recording took ∼30 min, with the animal in the setup for ∼180 min in total. Data was recorded at 20 kHz (PCIe-6321, National instruments and Intan board RHD2000 Evaluation Board version 1.0).

### *In vitro* electrophysiology

Mice were deeply anesthetised via intraperitoneal (i.p.) injection of ketamine (95 mg kg^-1^) and xylazine (4.5 mg kg^-1^), followed by transcardial perfusion with ice-cold, oxygenated (95 % O_2_, 5 % CO_2_) artificial cerebrospinal fluid (ACSF) containing (in mM): 118 NaCl, 2.5 KCl, 1.25 NaH_2_PO_4_, 1.5 MgSO_4_, 1 CaCl_2_, 10 Glucose, 3 Myo-inositol, 30 Sucrose, 30 NaHCO_3_; pH = 7.4. The brain was rapidly excised and coronal sections of 300 µm thickness containing the SC and PAG were cut using a Linear-Pro7 vibratome (Dosaka, Japan). Slices were left to recover for 20 min at 35°C, followed by a slow cool down to room temperature (RT) over 40 – 60 min. After recovery, one slice was transferred to the recording chamber (RC-26GLP, Warner Instruments, Holliston, MA, USA) and superfused with ACSF containing 2 mM CaCl_2_ and 20 µM bicuculline methiodide (Tocris) at a rate of 3 – 4 ml/min at RT. In experiments using 100 nM α-dendrotoxin (Alomone Labs, Jerusalem, Israel), 0.1 % bovine serum albumin (BSA) was added to the ACSF. Glass pipettes (B150-86-10, Sutter Instrument, Novato, CA, USA) with resistances of 3 – 4 MΩ were crafted using a P1000 horizontal pipette puller (Sutter Instrument) and filled with internal solution containing (in mM): 130 K-Gluconate, 10 KCl, 5 MgCl_2_, 5 MgATP, 0.2 NaGTP, 0.5 EGTA, 5 HEPES; pH 7.4 adjusted with KOH. Biocytin (0.2 – 0.3 %) was added to the internal solution for post hoc morphological reconstruction. Neurons of the dPAG or the dmSC were visually identified using an infrared differential interference contrast video system in a BX51 microscope (Olympus, Tokyo, Japan). Electrical signals were acquired at 20 – 50 kHz and filtered at 4 kHz using a Multiclamp 700B amplifier (Molecular Devices, San Jose, CA, USA) connected to a Digidata 1440A digitizer (Molecular Devices) with pClamp10 software (Molecular Devices). Spontaneous excitatory postsynaptic currents (sEPSCs) were recorded at –60 mV for at least 1 min. Intrinsic membrane properties and neuronal excitability were measured in current clamp mode by applying current steps from – 20 to 200 pA for 1 s in 10 pA increments. Access resistance was constantly monitored between protocols and recordings with access resistances exceeding 20 MΩ or with changes in access resistance or holding current by more than 20 % were discarded. After recordings, the pipette was carefully withdrawn and the slice was transferred to 4 % paraformaldehyde (PFA) in PBS solution.

### Proteomics

#### Samples Preparation

Six biological replicates from cortex, superior colliculus, and periaqueductal gray tissue were dissected from *Setd5^+/+^* and *Setd5^+/-^*adult mice, immediately frozen in liquid nitrogen, and conserved at -70°C until further processing. Tissues were Dounce homogenized 15 times in RIPA buffer (50 mM Tris-HCl pH 8, NaCl 150 mM, 0.2% sodium dodecyl sulfate, 0.5% p/p sodium deoxycholate, 1 % Triton X-100, 1x protease inhibitor cocktail (P8849 Roche), and 1 mM phenylmethylsulfonyl fluoride (PMSF)) and centrifuged at 16000g 4 °C for 20 min. Supernatants were further collected and stored at -20 °C. Total protein quantity was measured with the bicinchoninic acid assay (BCA) and a total amount of 30 µg of protein per sample was used for mass spectrometry measurements. All samples were cleaned up by SP3 using a commercial kit (PreOmics GmbH, 50 mg of beads per sample), then processed using the iST kit (PreOmics GmbH) according to the manufacturer’s instructions. Tryptic digestion was stopped after overnight incubation and cleaned-up samples were vacuum dried. Finally, samples were vacuum dried then re-dissolved with 10 min sonication in the iST kit’s LC LOAD buffer.

#### LC-MS/MS analysis

Samples were analysed by LC-MS/MS on a nanoElute 2 nano-HPLC (Bruker Daltonics) coupled with a timsTOF HT (Bruker Daltonics). Samples were first concentrated over an Acclaim PepMap trap column (5.0 µm C18-coated particles, 0.5 cm * 300 µm ID, ThermoFisher Scientific P/N 160454), then bound onto a PepSep XTREME column (1.5 µm C18-coated particles, 25 cm * 150 µm ID, Bruker P/N 1893476) heated at 50°C and eluted over the following 125 min gradient: solvent A, MS-grade H₂O + 0.1% formic acid; solvent B, 100% acetonitrile + 0.1 % formic acid; constant 0.60 nL/min flow; B percentage: 0 min, 2%; 100 min, 18 %; 125 min, 35 %, followed immediately by a 5 min plateau at 95 %. MS method: M/Z range = 350-1200 Th, ion mobility range = 0.7-1.4 1/K0; transfer time = 60 µs, pre-pulse storage time = 12 µs, ion polarity = positive, scan mode = dia-PASEF; TIMS parameters: ramp time = 100 ms, accumulation time = 100 ms; PASEF parameters: ms/ms scans = 10, total cycle time = 1.16721 s, charge range = 0-5, intensity threshold for scheduling = 2,500, scheduling target intensity = 20,000, exclusion release time = 0.4 min, reconsider precursor switch = on, current/previous intensity = 4, exclusion window mass width = 0.015 m/z, exclusion window v·s/cm² width = 0.015 V*s/cm².

Raw files were searched in DiaNN version 1.8.1 in library-free mode against a Mus musculus proteome obtained from UniProtKB. Match-Between-Runs was turned off. Fixed cysteine modification was set to Carbamidomethylation. Variable modifications were set to Oxidation (M), Acetyl (protein N-term) and Phospho (STY). Data was filtered at 1% FDR. DiaNN’s output was re-processed using in-house R scripts, starting from the main report table. MS1 intensities were re-normalized using the Levenberg-Marquardt procedure to minimise sample-to-sample differences. The long format main report table was consolidated into a wide format peptidoforms table, summing up quantitative values where necessary. Peptidoform intensity values were re-normalized using the Levenberg-Marquardt procedure and corrected using the ComBat function from the sva package to remove the replicate-related batch effect. Protein groups were inferred from observed peptides, and quantified using an in-house algorithm which: i) computes a mean protein-level profile across samples using individual, normalised peptidoform profiles (“relative quantitation” step), ii) following the best-flyer hypothesis, normalises this profile to the mean intensity level of the most intense peptidoform (“unscaled absolute quantitation” step); for protein groups with at least 3 unique peptidoforms, only unique ones were used. Estimated expression values were log10-converted and re-normalized using the Levenberg-Marquardt procedure. P-values were adjusted using the Benjamini-Hochberg (FDR) method. Average log10 expression values were tested for significance using a two-sided moderated t-test per samples group. Significance thresholds were calculated using the Benjamini-Hochberg procedure for False Discovery Rate values of 1%, 5%, 10 and 20%. For all tests, regulated protein groups were defined as those with a significant P-value and a log2 ratio greater than 5% of control-to-average-control ratios.

### Histology

For fluorescent immunostainings and histological confirmation of fibre and probe locations, adult *Setd5*^+/+^ and *Setd5*^+/-^ mice were transcardially perfused with 4 % paraformaldehyde (PFA) and their brains were dissected, post-fixed overnight in 4 % PFA, washed with 1X PBS, dehydrated in a 30 % sucrose solution and sectioned with a freezing microtome (SM2010R, Leica) to 100 μm (for probe/fibre/cannula confirmation) and 40 μm (for immunostainings) thick slices. For histological confirmation of probe and fibre locations, the sections were rinsed in PBS, stained with 4’,6-diamidino-2-phenylindole (DAPI; 2 mM in PBS) and mounted on Superfrost glass slides in a mounting medium containing 1,4-diazabicyclooctane (DABCO; Sigma, D27802) and Mowiol 4-88 (Sigma, 81381). For Kv1.1 immunostainings, tissue was rinsed and then blocked for 1 hour at room temperature in a buffer solution containing 10% normal donkey serum (Abcam, ab7475), 0.1 % Triton X-100 in PBS. Primary antibodies (Kv1.1, Rabbit, 1:200; Rb-Af400, Frontier Institute) were diluted in a blocking buffer and incubated for 48 hours at 4 oC. The tissue was then rinsed 3 times in PBS and left overnight to wash at 4 oC. Secondary antibodies (Alexa-594 Donkey anti-rabbit, 1:1000; R37119, Thermo Fisher) were diluted in 1X PBS and incubated for 2 hrs at room temperature in the dark, before being washed three more times in 1X PBS before being counter-stained with DAPI and mounted as for histological sections described above. Sections were imaged with either a confocal microscope (LSM800, Zeiss), at 63X or 20X magnification, or a wide-field microscope (VS120, Olympus), at 10X magnification.

For visualisation of the dPAG/dmSC cells recorded *in vitro* and the neurobiotin infused into the cannula implanted mice, tissue sections were washed in 1X PBS for 10 minutes then a solution of Streptavidin (1:100, 1 mg/ml, Thermo Fisher Scientific, S11227) in PBS with 0.3 % Triton-X was added and the samples were left at 4 °C overnight. The tissue was then washed three times with PBS, in vitro slices were mounted into a custom designed chamber for thick slices and imaged with a confocal microscope (LSM800, Zeiss) at 20X (90 μm projection, 2 μm sections) and 40X (50 μm projection, 1 μm sections).

### Western blot

Tissue samples from the superior colliculus, periaqueductal grey (PAG) and cortex were dissected from *Setd5^+/+^* (n = 3) and *Setd5^+/-^* (n = 3) mice. PAG tissue was dissected using a 2 mm diameter biopsy punch (ID: 2 mm, OD: 3 mm, Fine Science Tools, 18035-02). Each purification was performed independently three times and samples were stored at -80 °C until further processing. Samples were Dounce homogenized 15 times with 300 mL RIPA buffer (50 mM Tris-HCl, 150 mM NaCl, 0.5 % sodium deoxycholate, 1 % Triton-X, 0.2 % SDS, completeTM protease inhibitor cocktail (Roche) and and 1 mM phenylmethylsulfonyl fluoride (PMSF, Sigma)) using the loose pestle and sonicated for 10 seconds. The lysed samples were centrifuged for 20 min at 16000 *g* and the supernatant was stored at -80°C. Total protein concentration was determined using the BCA protein assay (Pierce). Proteins were boiled for 5 min in Laemmli sample buffer before being separated at 200 V for 40 min using TGX stain-free precast gels (Bio-Rad). Stain-free protein gels were activated with UV light for 5 min and then transferred to PVDF membranes (Bio-Rad) using a turbo transfer system (Bio-Rad). Membranes were blocked for 1 hr in TBST (Tris-buffered saline with 0.1 % Tween 20) supplemented with 5 % bovine serum albumin (Sigma) and then incubated overnight at 4 °C with anti-Kv1.1 primary antibody (Rb-Af400, Frontier Institute, 1:100). After primary antibody incubation, membranes were washed four times with TBST for 5 min and then incubated with anti-rabbit IRDye 800CW (LI-COR Biosciences, dilution 1:4000) for 1 hr at room temperature.

## QUANTIFICATION AND STATISTICAL ANALYSIS

### Analysis of behavioural data

For all behavioural experiments, the initial tracking and analysis of the escape behaviour were done blind through automatic analysis scripts and the genotypes were added later when pooling the data across animals. Automatic image processing of the behavioural video frames identified the central position and the heading direction of the mouse.

#### Standard looming escape response protocol

Looming escape responses (LER) were assessed by analysing a peri-stimulus time period starting from 3 s before the stimulus until 10 s after the stimulus onset. The speed of the animal was calculated as a moving mean over 83 ms time bins and responses were classified as escapes if the mouse returned to the shelter within 7 s after stimulus onset and reached a maximum speed of at least 20 cm s^-1^. Behavioural metrics were calculated by finding the mean value across all escape trials for each animal and then averaging across mice from each genotype. The maximum escape speed was calculated as the peak value of the speed trace between the onset of the escape and entry into the shelter. The reaction time of the mouse was calculated as the time from stimulus onset till the mouse reached the maximum speed during its escape back to the shelter. This reaction time was then used to group the responses of the animals depending on which loom presentation the peak of their escape response was recorded. The number of looms triggered is the total number of loom stimuli triggered across the 5 days of testing, not including the reward trial. To analyse the immediate change in speed following stimulus onset, trials were pooled across animals from each genotype. The difference between the average speed ±50 ms around the start of the stimulus (S_at_) and the average speed in the time 300-800 ms after the start of the stimulus (S_im_) was calculated separately for trials where the mouse initiated an escape within the presentation of the first looming stimulus or after the presentation of the first looming stimulus. Responses were categorised based on whether the S_im_ was higher (> 1 standard deviation (SD)), lower (<1 SD) or within 1 SD from the average speed at the time of triggering the stimulus (S_at_) across all mice from the same genotype and the distribution of trials in each category was analysed using a X^2^ test of independence.

#### Exploratory behaviour and positional controls

Exploratory behavioural metrics during the acclimatisation period were calculated from videos of the first 10 min the mice were placed within the arena. A shelter exit was defined as an episode where the full body of the mouse moved from the inside to the outside of the shelter. The maximum distance travelled during an exit was defined as the maximal distance reached in a straight line between the centre of the mouse and the centre of the shelter. The exit duration was defined as the time between when the full body of the mouse crossed over until it re-crossed the shelter boundary. The interbout interval during exploration was the time between subsequent shelter exits. The centre position of the mouse, [Xm, Ym], was used to quantify the percentage of the 10 min recorded that the mouse was either outside of the shelter, in the centre of the area (defined as a circular area, centred on the centre of the arena where the loom was presented, [X_c_, Y_c_], with a 7 cm radius) or at the edge of the area (defined as outside of a circular area, centred on [X_c_, Y_c_] with a 14 cm radius). To understand the effect of certain parameters of the mouse’s position and location within the arena on its LER response we calculated the distance of the mouse from shelter (defined as the straight line distance between [X_m_, Y_m_] and [X_s_, Y_s_]) and the heading angle of the mouse at the time of stimulus start (defined as the angle (B C) between point B, the tip of the mouse’s snout, A, the position between the two back paws of the mouse and C the centre of the shelter, [X_s_,Y_s_]). The position of the mouse’s body parts was located using DeepLabCut tracking[53].

#### Looming escape response protocol in the presence of a food reward

Food reward experiments were analysed with the same scripts as the standard LER experiments. When quantifying the reaction time and maximum speed only trials where the animals met the escape criteria were included. Four *Setd5*^+/-^ trials were removed from the final analysis because the mouse was interacting with the banana chip reward during the stimulus presentation. The number of looms and number of exits over the reward trials were calculated as the mean across animals from of the total per animal across genotypes.

#### Different contrast experiments

Trials were grouped by stimulus contrast and pooled across animals from the same genotype. A repeated measure ANOVA analysis followed by a Bonferroni correction for multiple analyses, was used to assess the effect of stimulus contrast on reaction time across genotypes and individual X^2^ tests of independence were used to assess the distribution of trials into the three response categories based on the change in immediate speed for each contrast. Only trials that fulfilled the escape criteria of having returned to the shelter within 7s of stimulus onset and reaching a maximum speed of >20 cm s^-1^ were included in the analysis.

#### In vivo cannula experiments with α-DTX

Trials were pooled across animals from each genotype and only trials that met the escape criteria used in the standard LER trials were included in the analysis of reaction time and maximum escape speed. Shelter exits were quantified as the average number of shelter exits per animal over the LER trials either with saline or with ɑ-DTX. Comparisons of *Setd5*^+/+^ or *Setd5*^+/-^ animals with saline versus ɑ-DTX were done with paired *t*-tests if the data reached the normality criteria, otherwise paired Wilcoxon’s rank sum tests were used. Unpaired Wilcoxon’s rank sum tests were used for comparisons across genotypes.

### Analysis of optogenetics experiments

Trials were grouped according to the laser power used and pooled across animals from each genotype. As in the analysis of the LER experiments, the speed of animal was calculated across an 83 ms time bin and the effect of laser stimulation was assessed by comparing the mean speed of the animal ±50 ms around the onset of the laser (S_at_) and 300-800 ms after the onset of the laser (S_im_). Trials from the 0.01 mW mm^-2^ condition were not subject to the escape criteria since laser stimulation at this intensity did not elicit an observed response, however, trials from all other laser power conditions had to meet the escape criteria used in the standard LER experiments in order to be included in further analyses. Trials were grouped by laser power intensity and S_at_ and S_im_ were compared for all trials by paired Wilcoxon signed-rank tests. Similar to the LER analysis, responses were individually grouped based on both the sign and the magnitude of the difference between S_at_ and S_im_, into three groups where S_im_ is either more than 1 standard deviation (SD) greater than S_at_, more than 1 SD less than S_at_ or within 1 SD of S_at_. Chi-Square tests were used to compare the distribution of the *Setd5*^+/+^ and *Setd5*^+/-^ trials at different laser powers.

For the control experiments, with GcaMP injected mice (Fig. S5) the Log_10_(**Δ**Speed) was calculated as the logarithmic of the ratio between S_im_ and S_at_. The change in Log_10_(**Δ**Speed) across laser powers was analysed using a two-way repeated measures ANOVA looking at the effect of genotype, laser power and their interaction, followed by a multiple comparisons analysis with Bonferroni correction.

### Analysis of *in vivo* silicon probe recordings

#### Preprocessing

Initial analysis was performed in MATLAB R2018b and later in MATLAB R2019a. Raw voltage traces were high-pass filtered (300 Hz) and spikes were detected and sorted using Kilosort2 (https://github.com/cortex-lab/Kilosort) and Phy as previously described[54], followed by manual curation. Firing rates histograms were calculated as the average firing rate across 16.7 ms time bins. Analog input channels (Photodiode, camera trigger) were extracted in MATLAB (MathWorks, Inc.) and arduino were recorded at a 20 kHz sample rate and were used for offline synchronisation. Custom-made scripts in MATLAB were used to extract the time points of the visual stimulus onsets and matched them to the corresponding video camera frame.

#### Determining the surface of the superficial superior colliculus (sSC)

Initial SC location was determined in real-time by auditory monitoring of multi-unit activity to a flashing stimulus and was later confirmed offline using a current source density analysis of the raw recording (see [31,55]) combined with histological reconstruction of the probe location visualised with DiI. CSD profiles were calculated for each recording and were used to align the depth of the units across recordings and across animals. The transition point between an overlying source and underlying sink was taken as the boundary between the stratum griseum superficiale (SGS) and the stratum opticum (SO). The boundary between the superficial (sSC) and intermediate/deep superior colliculus (idSC) was defined as 100 μm below this inflection depth (ID), taking into account the thickness of the SO. sSC was defined as 300 μm above the ID to 100 μm below the ID, and idSC was defined as the area below 100 μm below the ID. Only units identified as within the sSC and units with at least 2 spikes within the stimulus presentation were used for analysis. For all analyses spikes were sorted into peristimulus periods based on synchronisation with the stimulus frames and firing-rate histograms were calculated over 16.7 ms time bins.

#### Analysis of baseline firing properties

Responses to a full field grey screen during the first 60s of recording were taken to determine the baseline firing properties of the recorded cells. The baseline firing rate per animal was calculated as the mean firing rate across cells in the 1000 μm below the sSC surface.

#### Analysis of responses to flash stimulus

Responses to a single flash stimulus (0.5 s OFF; 1 s ON; 0.5 s OFF) were generated by averaging the firing-rate histograms across 10 repetitions of the stimulus. The responsiveness of single units was defined based on the p-value generated from running a ZETA test[56] on the average firing rate histograms using the 30 s period before the stimulus as baseline and the 2 s flash stimulus as the response time. Clustering of responsive units was done using K-means and the number of clusters was determined by identifying the elbow point when analysing the Calinski-Harabasz index. Clustering was done with units from both genotypes pooled together, then later split based on genotype. The proportion of flash-responsive cells for *Ptchd1* and *Cul3* animals were determined for each animal by dividing the number of sSC units that were p < 0.05 by the total number of sSC units found.

#### Analysis of responses to loom stimulus

Responses to a single loom stimulus were obtained by averaging the responses to the first loom of the loom bout stimulus (consisting of 5 consecutive looms) across 10 repetitions. Responsiveness was again defined as units having a *p*-value (determined using the ZETA test) of < 0.05, with the 30 s preceding the loom stimulus as baseline and the 0.7 s of first loom presentation as the response time. The average firing rate was calculated as the overall average of the mean firing rate recorded across each of the 5 loom presentations individually for 10 repetitions. Maximum firing was similarly calculated as the mean maximum firing during each of the 5 looms, averaged across 10 repetitions. The time-to-peak was calculated as the mean of the time to the peak firing within each of the 5 looms, averaged over the 10 repetitions.

#### Analysis of responses to shifting white noise stimulus

To further analyse the visual receptive fields (RFs) of the units recorded we analysed their responses to a shifting white noise stimulus[52] and calculated them as spike-triggered averages. The location of the centre of the receptive field was estimated by finding the pixel with the highest variance over a peri-spike time period and the final RF was cropped to 160 x 160 pixels centred on this pixel with highest variance. Signal-to-noise ratio (SNR) for each neuron is defined as *10*⋅*log_10_(max_power_/noise_power_)*, where *max_power_* is the square of the pixel value at the receptive field centre, and *noise_power_* the variance of all the values in a 5 pixels border around the edge of the RF. Visually-responsive neurons were defined as those with SNRs in the 80^th^ percentile of the SNRs of all sSC units or above. The centre of the RF was assigned as ON or OFF based on the sign of the mean of the central pixel at the centre of the receptive field. The proportion of OFF RFs was calculated as the ratio between visually-responsive cells with OFF RFs and the total number of visually-responsive cells in the sSC. The radius of the RF centre was calculated as the width at half-peak of the spatial RF profile. The average radius of the receptive fields of an animal was calculated using only the visually-responsive units. Temporal receptive fields were clustered similarly to flash responses, using K-means to cluster the responses and the number of clusters was determined using the Calinski-Harabasz index.

#### Analysis of responses to moving bars and gratings stimuli

For each direction the mean across trials was calculated and normalised to the baseline firing in the 30 s before stimulus onset. The preferred direction (D_p_) was defined as the direction with the greatest normalised response and the non-preferred direction (D_np_) was taken as 180 degrees from that angle. The direction selectivity index (DSI) was then computed as (D_p_-D_np_)/(D_p_+D_np_) with values falling in a range from 1 (high direction selectivity) to 0 (low direction selectivity). Units were defined as being ‘direction selective’ (DS) if their DSI was in the 80^th^ percentile or higher out of all units recorded within the sSC. The proportion of direction selective cells per animal was found by quantifying the proportion of DS sSC units divided by all sSC units recorded for that animal.

#### Analysis of behavioural data from head-fixed visual stimulation

The motion of the animal was determined by sorting the difference between pixels in neighbouring frames with K-means into two groups. The proportion of time each animal spent running was calculated as the proportion of frames that were registered as ‘moving’ over the total number of frames. The pupillary light reflex (PLR) of the animals was assessed by tracking the change in pupil size across the repetitions of the flash stimulus. A deep neural network approach for markerless tracking[53] was used to extract points around the pupil. The diameter of the pupil was then calculated by fitting an ellipse to these points. Since we observed, and it is known, that the size of the pupil is greatly affected by locomotion, when analysing the PLR we only included repetitions in which the mice were running. An animal was considered to be running during a repetition if the mouse was classified as ‘moving’ in over half of the video frames covering that repetition.

### Analysis of in vitro patch clamp electrophysiological recordings

Whole-cell recordings were analysed in Clampfit (Molecular Devices) and MiniAnalysis Program (Synaptosoft). Resting membrane potential (RMP) was measured as the average membrane potential prior to current-injection. Input resistance (R_in_) was calculated from the slope of the linear fit of the hyperpolarising current/voltage relationship. Membrane time constant τ was measured from the exponential fit of the membrane potential change in response to a –10 or –20 pA current-injection. Membrane capacitance (C_m_) was calculated from C_m_ = τ/R_in_. Action potential firing was quantified in Clampfit using the event detection function. Phase-plane plots were performed to investigate the dynamic changes in the membrane potential over time (dV/dt) and were generated using the spikes from the first current step necessary to elicit action potentials (rheobase current). Soma size was calculated by fitting an oval to the projected image (63X) of the neurons and calculating the area of that oval.

### Analysis of proteomics

MS files were searched in DiaNN version 1.8.1 in library-free mode against a *Mus musculus* proteome obtained from UniProtKB. Match-Between-Runs was turned off. Fixed cysteine modification was set to Carbamidomethylation. Variable modifications were set to Oxidation (M), Acetyl (protein N-term) and Phospho (STY). Data was filtered at 1% FDR. DiaNN’s output was re-processed using in-house R scripts, starting from the main report table. MS1 intensities were re-normalized using the Levenberg-Marquardt procedure to minimize sample-to-sample differences. The long format main report table was consolidated into a wide format peptidoforms table, summing up quantitative values where necessary. Peptidoform intensity values were re-normalized using the Levenberg-Marquardt procedure and corrected using the ComBat function from the sva package to remove the replicate-related batch effect. Protein groups were inferred from observed peptides, and quantified using an in-house algorithm which: *i)* computes a mean protein-level profile across samples using individual, normalized peptidoform profiles (“relative quantitation” step), *ii)* following the best-flyer hypothesis, normalizes this profile to the mean intensity level of the most intense peptidoform (“unscaled absolute quantitation” step); for protein groups with at least 3 unique peptidoforms, only unique ones were used, otherwise razor peptidoforms were also included; Phosphopeptidoforms and their unmodified counterparts were excluded from the calculations. Estimated expression values were log10-converted and re-normalised using the Levenberg-Marquardt procedure. P-values were adjusted using the Benjamini-Hochberg (FDR) method. Average log10 expression values were tested for significance using a two-sided moderated t-test per samples group and a global F-test (limma). Significance thresholds were calculated using the Benjamini-Hochberg procedure for False Discovery Rate values of 1 %, 5 %, 10 and 20 %. For all tests, regulated protein groups were defined as those with a significant P-value and a log2 ratio greater than 5 % of control-to-average-control ratios. GO terms enrichment analysis was performed comparing for each test regulated against observed protein groups (topGO). Statistical analysis was performed for Phospho-modified peptides as for protein groups, normalizing ratios to parent protein group(s).

### Analysis of histological data

For comparison between the Kv1.1 expression in *Setd5*^+/+^ and *Setd5*^+/-^ tissue 30 μm stacks were imaged with 1 μm sections at 63X magnification, or 55 μm stacks were imaged with 2 μm sections at 20X magnification, and the same laser power settings were used for all images at a given magnification.

### Analysis of western blot data

The Bio-Rad ChemiDoc MP system was used for imaging and acquisition and images were quantified using ImageJ software. Protein levels are displayed as fold change. Student’s *t* test was used to test for significance (MATLAB, MathWorks, Inc.). Data represent mean ± SEM obtained from three mice per genotype.

### Statistics

Analyses were performed in custom-written MATLAB (2019a) scripts (MathWorks, Inc.). Shapiro-Wilk and Levene’s tests were used to test the normality and the equal variances of the datasets, respectively and parametric or non-parametric tests were used accordingly. Two-tailed tests were used unless specified. All statistical tests are reported in the text and appropriate Fig. legends (^∗^*p* < 0.05, ^∗∗^*p* < 0.01, ^∗∗∗^*p* < 0.001). In bar plots, the mean ± s.e.m. are shown, unless otherwise stated. Boxplots display the median and IQR.

## Acknowledgments

We thank Prof. Peter Jonas for his suggestion on the involvement of potassium channels and members of the Neuroethology group for their comments on the manuscript. Katalin Szigeti and Julie Murmann for experimental help. This research was supported by the Scientific Service Units of IST Austria through resources provided by the Lab Support Facility, the

Imaging and Optics Facility, the Machine Shop Unit and the Preclinical Facility, especially Freyja Langer and Michael Schunn.

## Author contributions

LB and MJ designed the study. LB analysed all data with help from OS. LB performed the behavioural, optogenetic, ɑ-DTX cannula, and *in vivo* electrophysiology experiments. PK performed the *in vitro* experiments. LB, MJ and OS developed the behavioural system and analysis pipeline. TVZ, OS and MJ developed the *in vivo* electrophysiology setup and LB the analysis pipeline. TM performed the protein level analysis. XC, GN and TR generated the *Ptchd1* mouse line. GN and RS provided mouse lines, reagents and gave advice on the study design. LB and MJ wrote the manuscript with help of the other authors.

## Data and materials availability

Proteomics, behavioural data, *in vivo* electrophysiological silicon probe data and *in vitro* electrophysiological patch clamp data will be deposited and made available as of the date of publication. Microscopy data reported in this paper will be shared by the lead contact upon request.

