## Supplementary Figures for "Recurrent impairments in visual perception and place avoidance across autism models are causally linked in the haploinsufficiency model of intellectual disability *Setd5*"

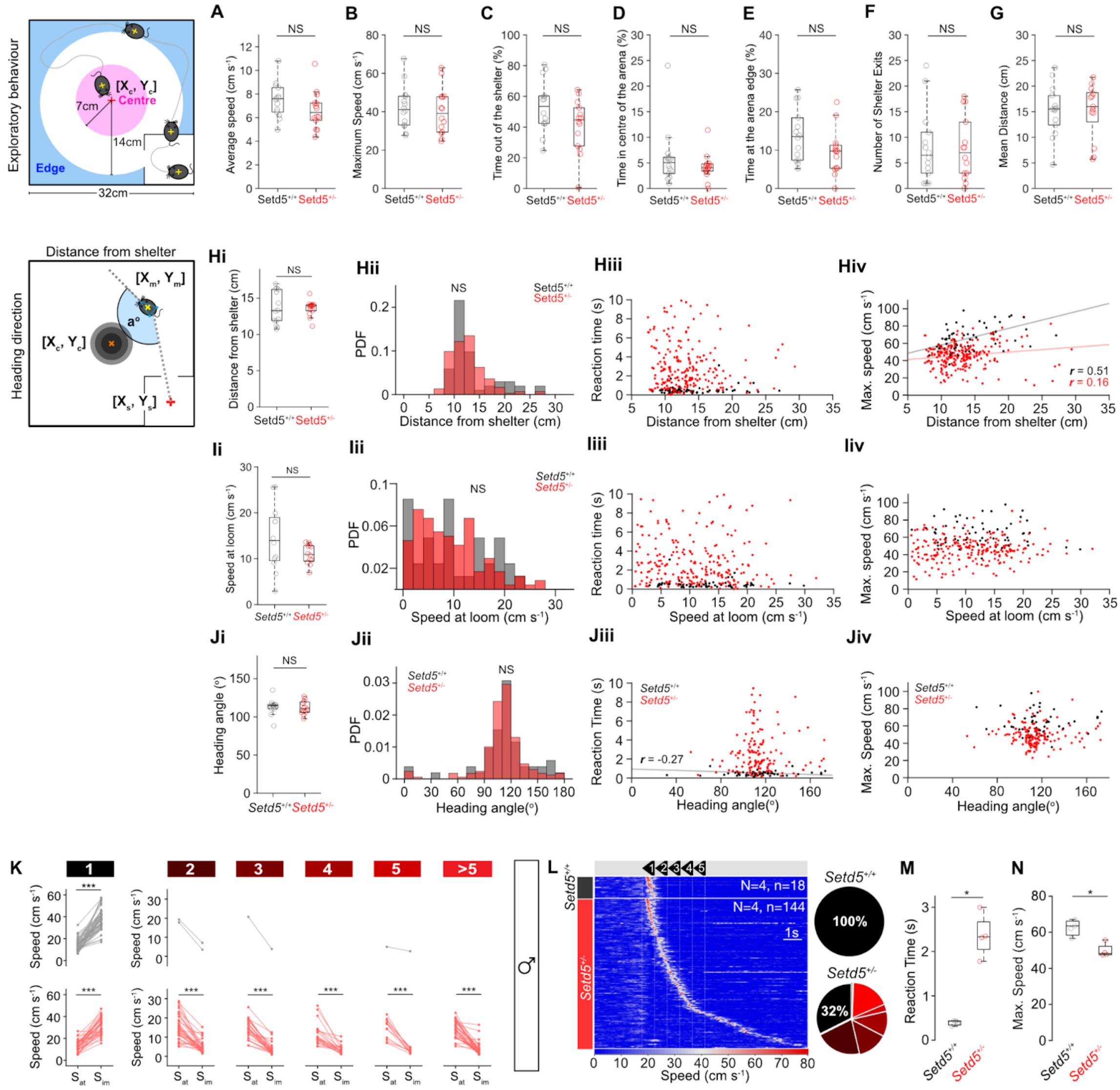


**Fig. S1. Additional *Setd5* behavioural controls.** No significant difference in the exploratory behaviour of the *Setd5*$^{+/+}$ and *Setd5*$^{+/-}$ animals during the pre-stimulus exploration. The top graphic depicts the dimensions and regions of the arena that are classified as the centre (pink, a circular region with a radius of 7cm from the centre of the arena [$X_c$,$Y_c$], and the edge (blue, the area outside of a circular area with a 14cm radius from [$X_c$, $Y_c$]. **a**, Average speed during pre-stimulus exploration (*Setd5*$^{+/+}$, 7.50 cms$^{-1}$; *Setd5*$^{+/-}$, 6.65 cms$^{-1}$, P = 0.151). **b**, Maximum speed (*Setd5*$^{+/+}$, 42.2 cms$^{-1}$; *Setd5*$^{+/-}$, 40.2 cms$^{-1}$, P = 0.642). **c**, Time spent out of the shelter (*Setd5*$^{+/+}$, 53.5 %; *Setd5*$^{+/-}$, 40.9 %, P = 0.062). **d**, Time spent in the centre of the arena (*Setd5*$^{+/+}$, 6.00 %; *Setd5*$^{+/-}$, 4.20 %, P = 0.448). **e**, Time spent at the edge of the arena (*Setd5*$^{+/+}$, 13.9 %; *Setd5*$^{+/-}$, 9.47 %, P = 0.076). **f**, Number of shelter exits (*Setd5*$^{+/+}$, 8.14; *Setd5*$^{+/-}$, 7.88, P > 0.999). **g**, Mean distance travelled during exit (*Setd5*$^{+/+}$, 15.2 cm; *Setd5*$^{+/-}$, 15.1 cm, P = 0.981). The lower graphic depicts the distance of the mouse from the shelter as the Euclidean distance between the centre of mouse, [$X_m$, $Y_m$] and the centre of shelter [$X_s$, $Y_s$], and the heading angle as the angle (a°) between a line drawn along the body axis of the mouse and another from the position between the back paws of the mouse and the centre of the shelter. The bottom graphic depicts how the directedness of the escape trajectory is calculated by finding the shortest Euclidean distance between the centre of the mouse and the edge of the shelter when the stimulus starts and the actual distance travelled between the stimulus start and the mouse re-entering the shelter. **hi**, Median distance from the shelter when the looming stimulus is triggered (*Setd5*$^{+/+}$, 13.2 cm; *Setd5*$^{+/-}$, 13.7 cm, P = 0.341). **hii**, Distribution of distances from the shelter across all trials, pooled by genotype ($p = 0.325$, two-way Kolmogorov-Smirnov test). **hiii**, Relationship between the distance to the shelter and the reaction time (*Setd5*$^{+/+}$, $p = 0.443$; *Setd5*$^{+/-}$, $p = 0.778$). **hiv**, Relationship between the distance to the shelter and the maximum escape speed reached during escape trials (black, *Setd5*$^{+/+}$, r = 0.51, $p < 0.001$; red, *Setd5*$^{+/-}$, r = 0.16, $p = 0.016$). **ii**, Median instantaneous speed when the looming stimulus is triggered (*Setd5*$^{+/+}$,15.1 cms$^{-1}$; *Setd5*$^{+/-}$, 12.0 cms$^{-1}$, P = 0.312). **iii**, Distribution of the instantaneous speeds across all trials, pooled by genotype ($p = 0.086$, two-way Kolmogorov-Smirnov test). **iiii**, Relationship between the instantaneous speed and the reaction time (*Setd5*$^{+/+}$, $p = 0.197$; *Setd5*$^{+/-}$, $p = 0.112$). **iiv**, Relationship between the instantaneous speed and the maximum escape speed reached during escape trials (*Setd5*$^{+/+}$, $p = 0.592$; *Setd5*$^{+/-}$, $p = 0.580$). **ji**, Median heading angle when the looming stimulus is triggered (*Setd5*$^{+/+}$,114°; *Setd5*$^{+/-}$, 111°, P = 0.194). **jii**, Distribution of heading angles across all trials, pooled by genotype ($p = 0.994$, two-way Kolmogorov-Smirnov test). **jiii**, Relationship between the instantaneous speed and the reaction time (*Setd5*$^{+/+}$, $r = -0.37$, $p = 0.014$; *Setd5*$^{+/-}$, $p = 0.333$). **jiv**, Relationship between the instantaneous speed and the maximum escape speed reached during escape trials (*Setd5*$^{+/+}$, $p = 0.635$; *Setd5*$^{+/-}$, $p = 0.188$). **k**, Significant increase in speed upon stimulus presentation when mice respond within the first loom presentation (*Setd5*$^{+/+}$, 57 trials, $p < 0.001$; *Setd5*$^{+/-}$, 38 trials, $p < 0.001$, paired *t*-test). $S_{at}$ is the mean speed of the animal ±50 ms of stimulus onset and $S_{im}$ is the mean speed of the animal 300-800ms after stimulus onset. Significant decrease in speed for *Setd5*$^{+/-}$ trials where the mice respond after the first loom (Within 2$^{nd}$ loom: 35 trials; within 3$^{rd}$ loom: 31 trials; within 4$^{th}$ loom: 31 trials; within 5$^{th}$ loom: 16 trials; after 5$^{th}$ loom: 45 trials, all $p < 0.001$, paired *t*-tests). **l**, Left, raster plot of mouse speed during escape trials for male *Setd5*$^{+/+}$ (top, n = 4, 18 trials) and *Setd5*$^{+/-}$ (bottom, n = 4, 144 trials) mice, sorted by reaction time. White, dotted vertical lines denote the start of each loom; white solid line denotes the end of the stimulus. Right, summary of the proportion of trials in which the mice respond *within each loom* for *Setd5*$^{+/+}$ (top) and *Setd5*$^{+/-}$ mice (bottom). Proportion of trials where mice escaped within the first loom: Setd5$^{+/+}$, 1.00, *Setd5*$^{+/-}$, 0.324, P = 0.028. Distribution of number of looms to escape, *Setd5*$^{+/+}$, 18 trials, *Setd5*$^{+/-}$, 144 trials, $p < 0.001$. **m**, Summary of the average reaction time per animal (Setd5$^{+/+}$, 18 trials, 0.394 s; *Setd5*$^{+/-}$, 130 trials, 2.10 s, P = 0.029). **n**, Summary of the average maximum escape speed per animal (Setd5$^{+/+}$, 18 trials, 63.3 cms$^{-1}$; *Setd5*$^{+/-}$, 130 trials, 49.9 cms$^{-1}$, P = 0.047). Box-and-whisker plots show median, IQR and range. Error Bars represent the SEM. P-values: Wilcoxon's test, p-values: Pearson's correlation, unless specified. Plotted linear fits depict the statistically significant correlations.

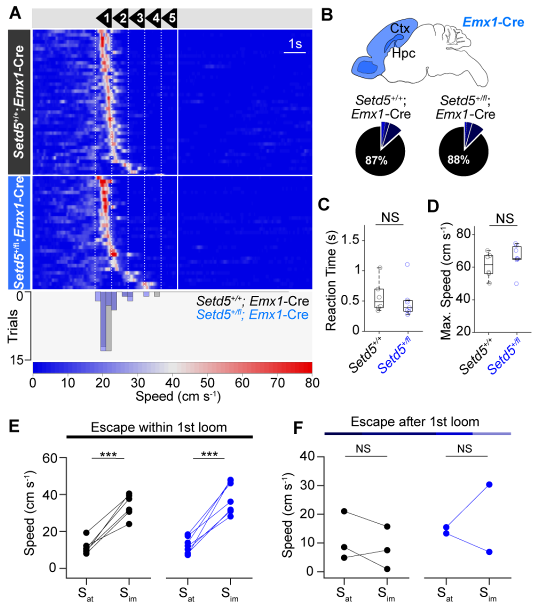


**Fig. S2. WT responses in cortical and hippocampal Setd5$^{+/-}$ animals. a**, Raster plot of mouse speed in response to the looming stimuli (white, dotted vertical lines denote the start of each loom; white solid line denotes the end of the stimulus) for *Setd5*$^{+/+}$; *Emx1*-Cre (top, n = 6, 42 trials) and *Setd5*$^{+/fl}$; *Emx1*-Cre (bottom, n = 6, 32 trials), sorted by reaction time. Bottom, Distribution of number of looms to escape across all trials (*Setd5*$^{+/+}$; *Emx1*-Cre, 32 trials, *Setd5*$^{+/fl}$;*Emx1*-Cre, 42 trials, p = 0.960, two-sample Kolmogorov-Smirnov (KS) test). **b**, Pictorial representation of the expression of *Emx1* in a sagittal section of a mouse brain, showing the localisation of the expression to the cortex and hippocampus (blue, top). Proportion of trials in which the mice respond within each loom for *Setd5*$^{+/+}$; *Emx1*-Cre (middle) and *Setd5*$^{+/fl}$; *Emx1*-Cre mice (bottom). **c**, Reaction times and (**d**) maximum escape speed (*Setd5*$^{+/+}$; *Emx1*-Cre, n = 6, 61.1 cms$^{-1}$; *Setd5*$^{+/fl}$;*Emx1*-Cre, n = 6, 66.4 cms$^{-1}$, P = 0.295). **e**, Speed immediately following the stimulus presentation for trials where the mice escape within the first loom presentation (left, *Setd5*$^{+/+}$; *Emx1*-Cre*,* n = 6, p < 0.001; right, *Setd5*$^{+/fl}$; *Emx1*-Cre, n = 6, p < 0.001). **f**, Speed immediately following the stimulus presentation for trials where the mice escape after the first loom presentation (left, *Setd5*$^{+/+}$*,* n = 3, p = 0.6248, paired *t*-test; right, *Setd5*$^{+/-}$, n = 2, p = 0.7546). P-values: Wilcoxon's ranked-sum test. p-values: two-tailed paired *t*-test, unless specified.

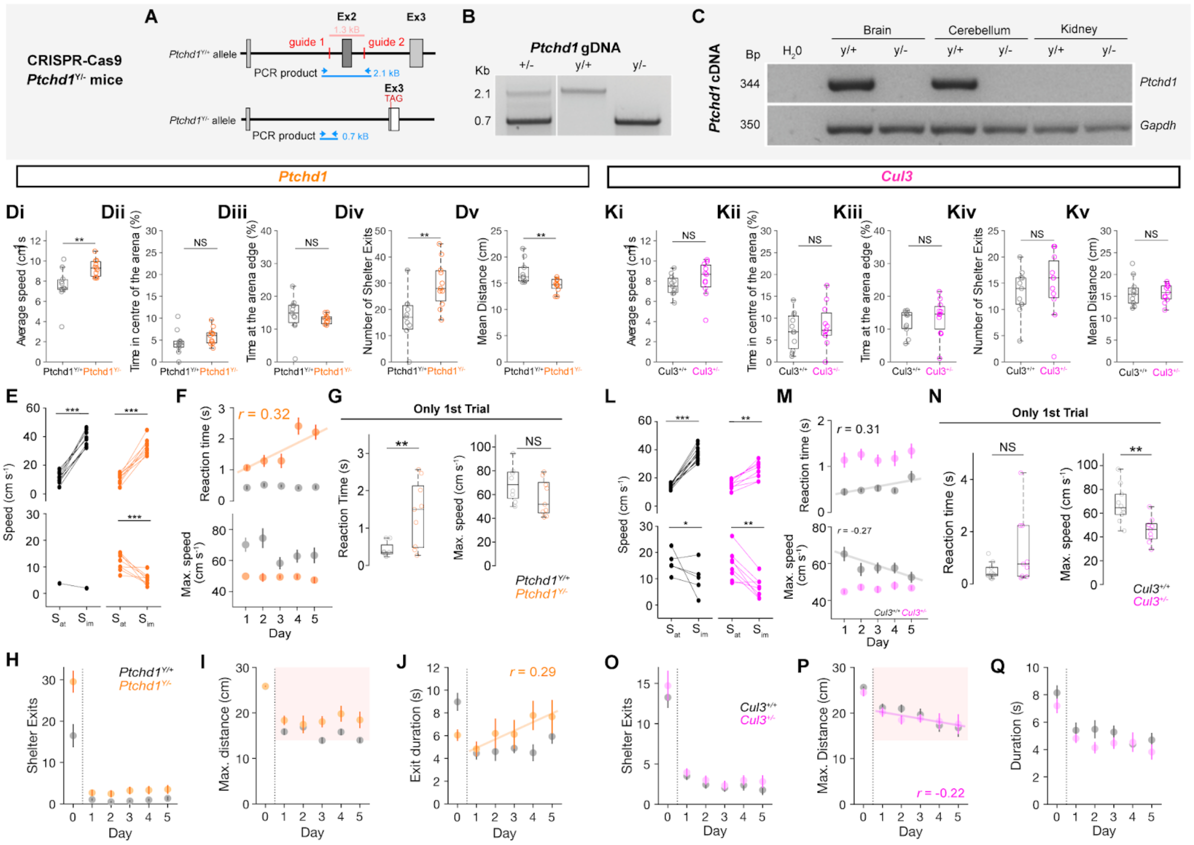


**Fig. S3. Ptchd1 animal model generation & Ptchd1 and Cul3 behavioural characterisation**. **a**, CRISPR-Cas9 design for the generation of *Ptchd1*$^{Y/-}$ mice. **b**, gDNA confirmation of the excised DNA (2.1 Kb in *Ptchd1*$^{Y/+}$ and 0.7 Kb in *Ptchd1*$^{Y/-}$). **c**, Confirmation of loss of Ptchd1 cDNA in different tissues in *Ptchd1*$^{Y/-}$ mice, and control *Gapdh* presence throughout. **d,** Exploration controls for *Ptchd1*. **di**, Average speed during pre-stimulus exposure acclimatisation (Ptchd1$^{Y/+}$, 7.53 cms$^{-1}$; Ptchd1$^{Y/-}$, 9.31 cms$^{-1}$, p = 0.003). **dii**, Time spent in the centre of the arena (Ptchd1$^{Y/+}$, 4.46 %; Ptchd1$^{Y/-}$, 5.86 %, p = 0.170). **diii**, Time spent at the edge of the arena (Ptchd1$^{Y/+}$, 14.3 %; Ptchd1$^{Y/-}$, 13.0 %, p = 0.148). **div**, Number of shelter exits (*Ptchd1*$^{Y/+}$, 17.0, *Ptchd1*$^{Y/-}$, 29.5, p = 0.003). **dv**, Average distance travelled during exit (*Ptchd1*$^{Y/+}$, 17.1 cm, *Ptchd1*$^{Y/-}$, 14.6 cm, p = 0.004). **e**, Speed change immediately following the stimulus presentation for trials where the mice escape within the first loom presentation (top left, *Ptchd1*$^{Y/+}$, n = 9, black, p < 0.001, two-tailed *t*-test; top right, *Ptchd1*$^{Y/-}$, n = 9, red, p < 0.001, two-tailed *t*-test). Speed change immediately following the stimulus presentation for trials where the mice escape after the first loom presentation (bottom left, *Ptchd1*$^{Y/+}$, n = 1, black; bottom right, *Ptchd1*$^{Y/-}$, n = 8, red, p < 0.001, two-tailed *t*-test). $S_{at}$ is the mean speed of the animal ±50 ms of stimulus onset and $S_{im}$ is the mean speed of the animal 300-800ms after stimulus onset. **f**, Top, relationship between reaction time and test day (*Ptchd1*$^{Y/+}$, $r = 0.029$, $p = 0.840$; *Ptchd1*$^{Y/-}$, $r = 0.317$, $p < 0.001$) and bottom, maximum escape speed and test day (*Ptchd1*$^{Y/+}$, $r = -0.254$, p = 0.07; *Ptchd1*$^{Y/-}$, $r = -0.040$, p = 0.556). **g**, Mean reaction time (left) and maximum escape speed (right) per animal, for the very first loom presentation. (Reaction time; *Ptchd1*$^{Y/+}$, 0.423 s, *Ptchd1*$^{Y/-}$, 1.39 s, P = 0.014. Maximum escape speed; *Ptchd1*$^{Y/+}$, 69.1 cms$^{-1}$, *Ptchd1*$^{Y/-}$, 57.3 cms$^{-1}$, P = 0.139). **h-j**, Relationship between test day and the number of exits from shelter (**h**, *Ptchd1*$^{Y/+}$, $r = 0.122$, p = 0.134; *Ptchd1*$^{Y/+}$, $r = 0.101$, p = 0.1936), the maximum distance travelled (**i**, *Ptchd1*$^{Y/+}$, $r = -0.213$, p = 0.071; *Ptchd1*$^{Y/+}$, 112 trials, $r = 0.103$, p = 0.278) and the duration of time spent outside of the shelter (**j**, *Ptchd1*$^{Y/+}$, $r = 0.148$, p = 0.214; *Ptchd1*$^{Y/+}$, $r = 0.286$, p = 0.002)**. k**, Exploration controls for *Cul3*. **ki**, Average speed during pre-stimulus exposure acclimatisation. Cul3$^{+/+}$, 7.60 cms$^{-1}$;

Cul3$^{+/-}$, 8.23 cms$^{-1}$, p = 0.3055. **kii**, Time spent in the centre of the arena (Cul3$^{+/+}$, 7.10 %; Cul3$^{+/-}$, 8.32 %, p = 0.532). **kiii**, Time spent at the edge of the arena (Cul3$^{+/+}$, 12.2 %; Cul3$^{+/-}$, 13.6 %, p = 0.480). **kiv**, Number of shelter exits (Cul3$^{+/+}$, 13.3, Cul3$^{+/-}$, 14.7, p = 0.526). **kv**, Average distance travelled during exit (Cul3$^{+/+}$, 15.8 cm; Cul3$^{+/-}$, 15.7 cm, p = 0.945). **l**, Top, speed change immediately following the stimulus presentation for trials where the mice escape within the first loom presentation (top left, *Cul3*$^{+/+}$, n = 10, black, $p < 0.001$, two-tailed t-test; top right, *Cul3*$^{+/-}$, n = 9, pink, p = 0.008, two-tailed t-test). Speed change immediately following the stimulus presentation for trials where the mice escape after the first loom presentation (bottom left, *Cul3*$^{+/+}$, n = 5, black, p = 0.017; bottom right, *Cul3*$^{+/-}$, n = 8, pink, p = 0.001, two-tailed t-test). **m**, Top, relationship between reaction time and test day (*Cul3*$^{+/+}$, $r$ = 0.3146, p = 0.012; *Cul3*$^{+/-}$, $r$ = 0.109, p = 0.233); bottom, relationship between maximum escape speed and test day (*Cul3*$^{+/+}$, $r$ = -0.2679, p = 0.033; *Cul3*$^{+/-}$, $r$ = 0.125, p = 0.1711). **n**, Mean reaction time (left) and maximum escape speed (right) per animal, for the very first loom presentation (Reaction time; *Cul3*$^{+/+}$, 0.446 s, *Cul3*$^{+/-}$, 1.26 s, P = 0.117. Maximum escape speed; *Cul3*$^{+/+}$, 67.5 cms$^{-1}$, *Cul3*$^{+/-}$, 45.8 cms$^{-1}$, P = 0.003). **o-p**, Relationship between test day and the number of exits from shelter (**o**, *Cul3*$^{+/+}$, $r$ = 0.083, p = 0.289; *Cul3*$^{+/-}$, $r$ = -0.1198, p = 0.1253), the maximum distance travelled (**p**, *Cul3*$^{+/+}$, $r$ = -0.081, p = 0.480; *Cul3*$^{+/-}$, $r$ = -0.224, p = 0.034) and the duration of time spent outside of the shelter (**q**, *Cul3*$^{+/+}$, $r$ = -0.036, p = 0.754; *Cul3*$^{+/-}$, $r$ = -0.091, p = 0.393). Box-and-whisker plots show median, IQR and range. p-values: Wilcoxon's test, P-values: Pearson's correlation analysis, unless specified. Trend lines are only drawn for significant correlations.

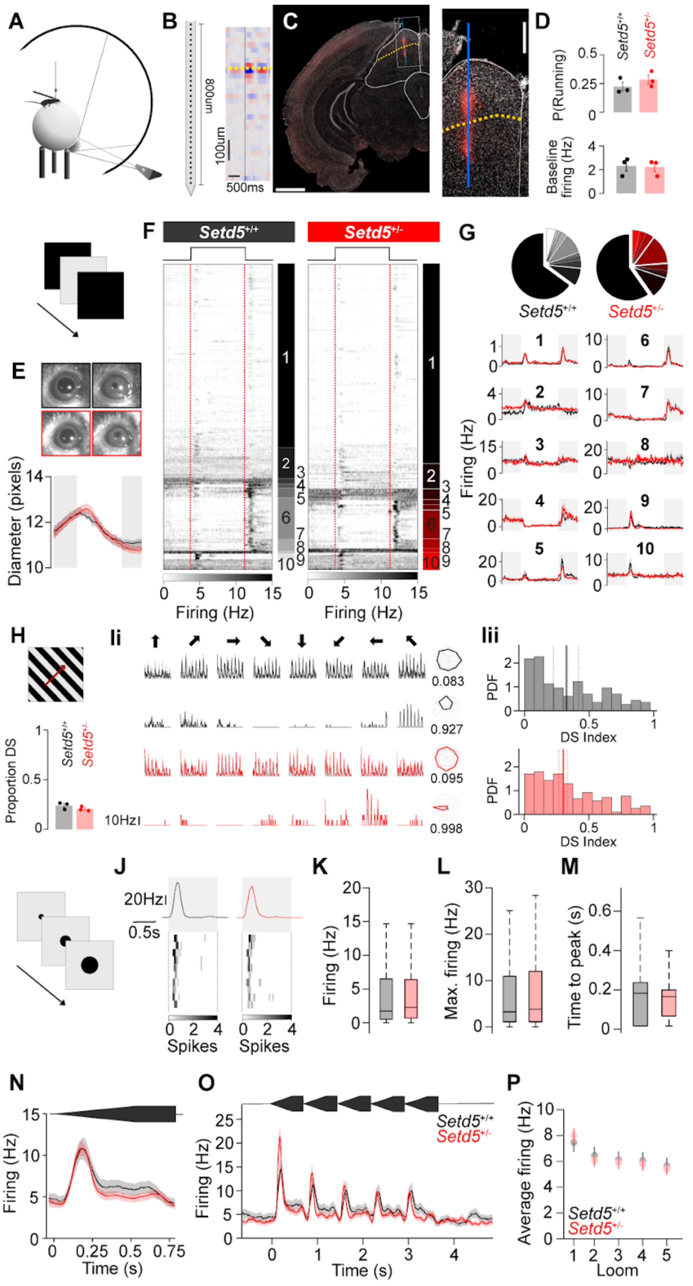


**Figure S4. Intact visual processing across the superior colliculus. a**, Schematic of the *in vivo* recording setup displaying the spherical treadmill and light projector illuminating the spherical screen. **b**, Left, Schematic of the 32-channel silicon probe used to record extracellular activity aligned with the current source density analysis of a single flash stimulus averaged over 10 presentations. Black vertical line represents the stimulus onset, yellow dotted line marks the inflection depth separating the current source and sink. **c**, Histological reconstruction of the probe position marked with DiI (120 µm coronal section, scale bar: 1 mm) with a close-up (scale bar: 250 µm). Yellow dotted line as in (**b**). **d**, Proportion of time the mice were moving during the recording sessions (P = 0.400, top) and SC's baseline firing rate (P = 0.880, bottom). **e**, Pupil dilation in response to the full-field flash stimulus. Example images of dilated and constricted pupils in responses to the OFF (left) and ON (right) periods of the flash. Setd5$^{+/+}$ (black), Setd5$^{+/-}$ (red) (P = 0.060). **f**, Sorted raster plot of neural responses to a single flash stimulus. Vertical red dotted lines indicate the on and offset of the flash stimulus. **g**, Proportion of cells in each cluster (top). Mean ± s.e.m. of the flash responses in each cluster (traces). **h**, Proportion of direction-selective SC cells (see Methods) to full field gratings (*Setd5*$^{+/+}$, 87 of 153 units, 0.287; *Setd5*$^{+/-}$, 159 of 259 units, 0.2432, P = 0.322, Wilcoxon's test). **i_i**, Firing rate of example units in response to full field gratings moving in 8 different directions for *Setd5*$^{+/+}$ (black) and *Setd5*$^{+/-}$ (red) with a summary polar plot of their direction selectivity (right) and the corresponding direction selectivity index (DSI) value. Bold lines show the mean response across 3 repetitions (light grey lines). **i_ii**, Distribution of DSI across SC units (top, *Setd5*$^{+/+}$, 153 units, median DSI = 0.323; bottom, *Setd5*$^{+/-}$, 259 units, median DSI = 0.3018, P = 0.7054, two-sample Kolmogorov-Smirnov test). **j**, Mean and spike raster plots from single Setd5$^{+/+}$ (black) and Setd5$^{+/-}$ (red) units to 10 repetitions of a single loom stimulus. **k-m**, Summary of mean, maximum and time-to-peak firing (**k**, P = 0.215; **l**, P = 0.100; **m**, P = 0.372). **n**, Mean ± s.e.m. response of all and (**o**) of the 5 consecutive loom stimuli for Setd5$^{+/+}$ (black) and Setd5$^{+/-}$ (red) units. **p**, Average firing to a single loom across the 5-loom stimulus (p = 0.732). Box-and-whisker plots show median, IQR and range. P-values: Wilcoxon’s test, p-values: two-way repeated measures ANOVA, unless specified.

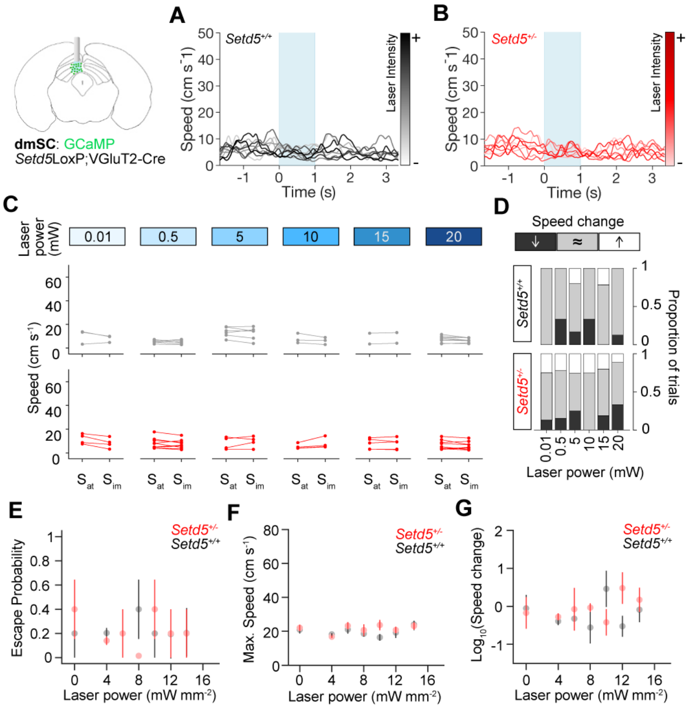


**Fig. S5. Optogenetic controls. a-b**, Mean speed responses to light stimulation at increasing laser intensities in *Setd5*$^{+/+}$;VGluT2-Cre (**a**) and *Setd5*$^{+/-}$;VGluT2-Cre (**b**) mice injected with AAV-GCaMP6m. Blue shaded areas show 1 s of 10 Hz light stimulation. **c**, Change in speed upon light activation at different laser intensities for *Setd5*$^{+/+}$ (top, n = 1. 0.01mW mm$^{-2}$: P = 0.700, 3 trials; 0.5 mW mm$^{-2}$: P = 0.472, 10 trials; 5.0 mW mm$^{-2}$: P > 0.995, 5 trials; 10.0 mW mm$^{-2}$: P = 0.700, 3 trials; 15.0 mW mm$^{-2}$: P = 0.667, 2 trials; 20.0 mW mm$^{-2}$: P = 0.442, 8 trials) and *Setd5*$^{+/-}$ (bottom, n = 1. 0.01mW mm$^{-2}$: P = 0.343, 4 trials; 0.5 mW mm$^{-2}$: P = 0.796, 9 trials; 5.0 mW mm$^{-2}$: P = 0.279, 4 trials; 10.0 mW mm$^{-2}$: P = 0.200, 4 trials; 15.0 mW mm$^{-2}$: P = 0.887, 4 trials; 20.0 mW mm$^{-2}$: P = 0.678, 10 trials) trials. $S_{at}$ is the mean speed of the animal ±50 ms of laser onset and $S_{im}$ is the mean speed of the animal 300-800ms after laser onset. **d**, Proportion of trials where the speed of the mouse increases (white: $S_{im} > S_{at}$ by more than 1 standard deviation (SD)), decreases (black: $S_{im} < S_{at}$ by more than 1 standard deviation (SD)) or doesn't change (grey: $S_{im}$ less than 1 SD different from $S_{at}$). 0.01mW mm$^{-2}$: 7 trials, p = 0.165; 0.5mW mm$^{-2}$: 19 trials, p = 0.624; 5mW mm$^{-2}$: 9 trials, p = 0.852; 10mW mm$^{-2}$: 7 trials, p = 0.766; 15mW mm$^{-2}$: 6 trials, p = 0.349; 20mW mm$^{-2}$: 18 trials, p = 0.815, $X^2$ test of independence. **e**, No relationship between escape probability and laser power (*Setd5*$^{+/+}$, *r* = -0.005, p = 0.955; *Setd5*$^{+/-}$, *r* = -0.047, p = 0.577). **f**, No relationship between maximum speed and laser power (*Setd5*$^{+/+}$, *r* = -0.014, p = 0.871; *Setd5*$^{+/-}$, *r* = 0.011, p = 0.8986). **g**, No relationship between the $Log_{10}$(Immediate ΔSpeed) and laser power (*Setd5*$^{+/+}$, *r* = -0.057, p = 0.496; *Setd5*$^{+/-}$, *r* = 0.283, p = 0.149). p-values: Pearson's correlation, P-values: paired Wilcoxon's tests, unless specified. *Setd5*$^{+/+}$; VGluT2-Cre, n = 1, 147 trials; *Setd5*$^{+/-}$; VGluT2-Cre, n = 1, 145 trials.

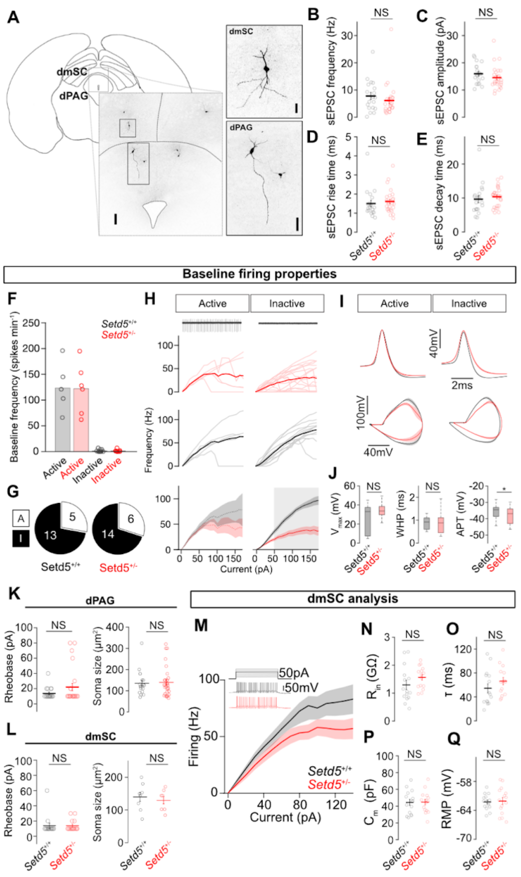


**Fig. S6. Supporting *ex vivo* slice electrophysiological analysis. a,** Location of dPAG and dmSC cells that were recorded and filled with biocytin (scale bar: 100 µm). Insets show close-ups of the biocytin-filled dPAG cells (top, scale bar: 50 µm) and dmSC cells (bottom, scale bar: 20 µm). **b-e**, Spontaneous neurotransmission in dPAG (*Setd5*$^{+/+}$: n = 6, 19 cells; *Setd5*$^{+/-}$: n = 7, 21 cells). **b**, Average sEPSC frequency (*Setd5*$^{+/+}$: 7.8 Hz; *Setd5*$^{+/-}$: 5.0Hz, P = 0.7743); **c**, average sEPSC amplitude (*Setd5*$^{+/+}$: 16.0 pA; *Setd5*$^{+/-}$: 14.0 pA, P = 0.704); **d**, average sEPSC rise time (*Setd5*$^{+/+}$: 1.50 ms; *Setd5*$^{+/-}$: 1.60 ms, P = 0.948); **e**, average sEPSC decay time (*Setd5*$^{+/+}$: 9.64 ms; *Setd5*$^{+/-}$: 10.5 ms, P = 0.838). **f**, Firing frequency at baseline splits the recorded cells into two groups. (Average baseline frequency: *Setd5*$^{+/+}$ high frequency, 5 cells, 123.5 Hz; *Setd5*$^{+/-}$ high frequency, 6 cells, 122.2 Hz; *Setd5*$^{+/+}$ low frequency, 13 cells, 1.12 Hz; *Setd5*$^{+/-}$ low frequency, 14 cells, 1.00 Hz). **g**, Proportion of recorded cells from *Setd5*$^{+/+}$ (left) and *Setd5*$^{+/-}$ (right) mice that exhibit high frequency (active) or inactive at baseline. **h**, Top, Example traces from an active (high frequency firing), putative GABAergic, cell (left) and an inactive, putative glutamatergic, cell (right) at baseline. Centre top, mean current-firing relationship of all putative GABAergic (left) and putative glutamatergic (right) *Setd5*$^{+/-}$ cells in red. Light pink lines represent individual cells. Centre bottom, mean current-firing relationship of all putative GABAergic (left) and putative glutamatergic (right) *Setd5*$^{+/+}$ cells in black. Light grey lines represent individual cells. Bottom, summary of the relationship between current injection and action potential firing for putative GABAergic (left, P = 0.560 for the effect of genotype) and glutamatergic (right, P < 0.001 for the effect of genotype) cells. The grey area indicates the current injection values that are significantly different between *Setd5*$^{+/+}$ cells (black) and *Setd5*$^{+/-}$ cells (red) found by a multiple comparisons analysis with Tukey correction. **i**, Average shape (top) and phase plane analysis (bottom) of the action potentials generated in the rheobase sweep of putative GABAergic (*Setd5*$^{+/+}$, 5 cells, 66 spikes; *Setd5*$^{+/-}$, 6 cells, 55 spikes) and putative glutamatergic (*Setd5*$^{+/+}$, 13 cells, 42 spikes; *Setd5*$^{+/-}$, 14 cells, 72 spikes) cells. **j**, Action potential kinetics analysis of putative GABAergic cells. Left to right: action potential amplitude ($V_{max}$) (*Setd5*$^{+/+}$, 66 spikes, 27.1 mV; *Setd5*$^{+/-}$, 55 spikes, 33.3 mV, p = 0.091), width at half-peak (*Setd5*$^{+/+}$, 66 spikes, 0.877 ms; *Setd5*$^{+/-}$, 55 spikes, 0.890 ms, p = 0.447); action potential threshold (*Setd5*$^{+/+}$, 66 spikes, -35.7 mV; *Setd5*$^{+/-}$, 55 spikes, -37.8 mV, p = 0.009). **k**, Left, rheobase current (*Setd5*$^{+/+}$: 19 dPAG cells, 13.7pA; *Setd5*$^{+/-}$: 21 dPAG cells, 22.1 pA, P = 0.609). Right, soma size (*Setd5*$^{+/+}$: 19 dPAG cells, 135.3µm; *Setd5*$^{+/-}$: 21 dPAG cells, 139.6 µm, P = 0.899, two-tailed *t*-test.) **l**, Left, rheobase current (*Setd5*$^{+/+}$: 15 dmSC cells, 14.0pA; *Setd5*$^{+/-}$: 15 dmSC cells, 14.0pA, P > 0.999, two-tailed *t*-test). Right, soma size (*Setd5*$^{+/+}$: 9 dmSC cells, 139.9µm; *Setd5*$^{+/-}$: 9 dmSC cells, 129.6 µm, P = 0.609, two-tailed *t*-test). **m-q**, Intrinsic properties of dmSC cells (*Setd5*$^{+/+}$: n = 4, 15 cells; *Setd5*$^{+/-}$: n = 4, 15 cells). **m**, Summary of the relationship between current-injection and action potential firing in dmSC cells (*Setd5*$^{+/+}$, black, n = 15; *Setd5*$^{+/-}$, red, n = 15, P=0.066 for a main effect of genotype, two-way repeated measures ANOVA). **n**, Input resistance (*Setd5*$^{+/+}$: 1.29 GΩ; *Setd5*$^{+/-}$: 1.56 GΩ, P = 1721). **o**, Tau (*Setd5*$^{+/+}$: 54.5 ms; *Setd5*$^{+/-}$: 66.4 ms, P = 0.229). **p**, Membrane capacitance (*Setd5*$^{+/+}$: 44.4 pF; *Setd5*$^{+/-}$: 44.7 pF, P = 0.961). **q**, Resting membrane potential (*Setd5*$^{+/+}$: -62.3 mV; *Setd5*$^{+/-}$: -62.0 mV, P = 0.795). p-values: Wilcoxon's tests, P values: repeated-measures analysis of variance (ANOVA), unless specified. In **h, i** and **m**, shaded areas represent s.e.m..

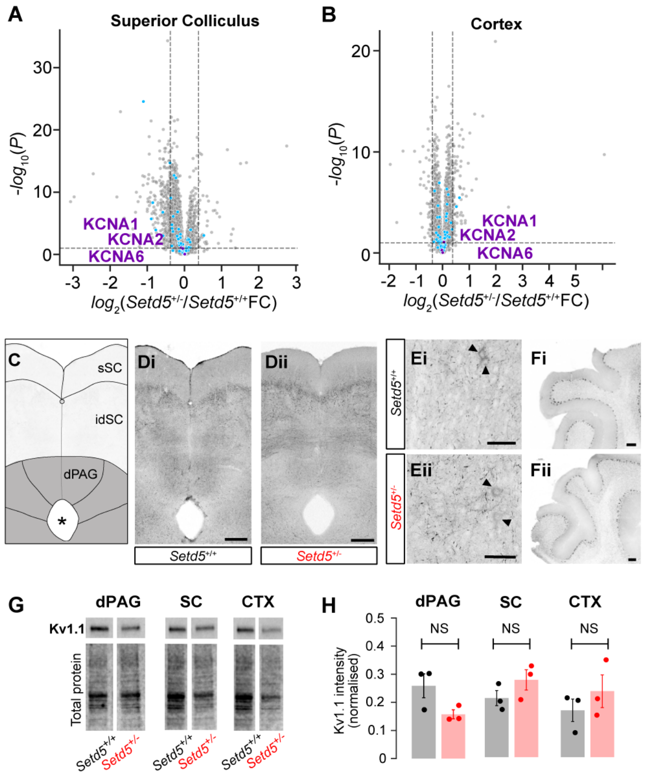


**Fig S7. Kv protein level analysis. a-b,** Volcano plots for differential protein levels in the superior colliculus (SC) and cortex (CTX) between adult Setd5$^{+/+}$ and Setd5$^{+/-}$ mice (n = 6 independent samples per genotype and brain area, cyan dots represent annotated proteins as ion-channels, and purple dots represent Kv1.1, Kv1.2 and Kv1.6, horizontal dashed line represents the significance threshold (p-value < 0.1 - two-sided moderated t-test), vertical dashed lines indicate fold change value of 0.4 between *Setd5*$^{+/+}$ and *Setd5*$^{+/}$. **c**, Schematic of SC and PAG regions of interest. **d-f**, Antibody staining for Kv1.1 in Setd5$^{+/+}$ (**di**, 55 µm projection; **ei**, 30 µm projection, **fi**, 100 µm projection) and Setd5$^{+/-}$ (**dii**, 55 µm projection; **eii**, 30 µm projection, **fi**, 100 µm projection). Arrowheads indicate somas stained for Kv1.1. **g**, Tissue specific western blots of Kv1.1 protein content in the dorsal periaqueductal grey (dPAG), the SC and the CTX for *Setd5*$^{+/+}$ (n = 3) and *Setd5*$^{+/-}$ mice (n = 3) and (**h**) their quantification (dPAG: *Setd5*$^{+/+}$, 0.259; *Setd5*$^{+/-}$, 0.157, P = 0.090. SC: *Setd5*$^{+/+}$, 0.210; *Setd5*$^{+/-}$, 0.275, P = 0.226. CTX: *Setd5*$^{+/+}$, 0.167; *Setd5*$^{+/-}$, 0.235, P = 0.392). Scale bar: **d:** 200 µm; **e**: 50 µm; **f**: 100 µm. P values are two-tailed Wilcoxon's signed-rank test.

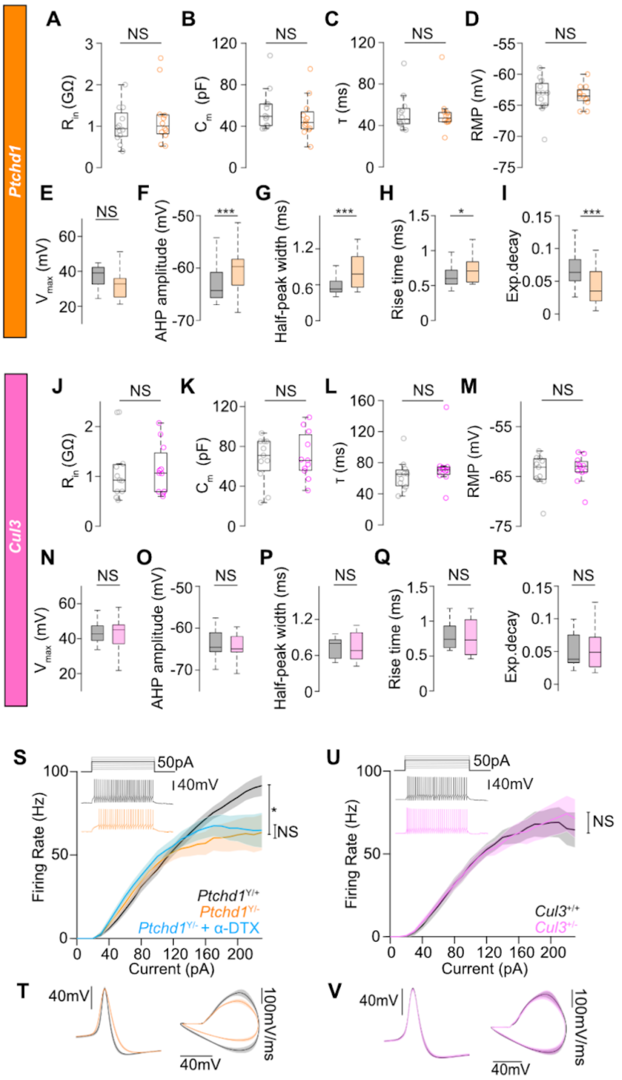


**Fig. S8. Electrophysiological *Ptchd1* and *Cul3* characterisation**. Intrinsic properties of dPAG cells in *Cul3* and *Ptchd1* animals. (**a,j**), Input resistance (**a**, *Ptchd1*$^{Y/+}$, 1.03 GΩ; *Ptchd1*$^{Y/-}$, 1.20 GΩ, p = 0.623; **j**, *Cul3*$^{+/+}$, 1.12 GΩ; *Cul3*$^{+/-}$, 1.14 GΩ, p = 0.896). (**b,k**) Membrane capacitance (**b**, *Ptchd1*$^{Y/+}$, 55.6 pF; *Ptchd1*$^{Y/-}$, 48.2 pF, p = 0.299; **k**, *Cul3*$^{+/+}$, 66.6 pF; *Cul3*$^{+/-}$, 70.8 pF, p = 0.896). (**c,l**), Membrane constant tau (**c**, *Ptchd1*$^{Y/+}$, 52.0 ms; *Ptchd1*$^{Y/-}$, 51.2 ms, p = 0.795; **l**, *Cul3*$^{+/+}$, 63.9 ms; *Cul3*$^{+/-}$, 74.3 ms, p = 0.212). (**d,m**), Resting membrane potential (**d**,*Ptchd1*$^{Y/+}$, -63.5 mV; *Ptchd1*$^{Y/-}$, -63.5 mV, p = 0.749; **m**, *Cul3*$^{+/+}$, -64.0 mV; *Cul3*$^{+/-}$, -63.5 mV, p = 0.844). Action potential kinetics for all spikes generated in the rheobase sweep. (**e, n**), Action potential amplitude ($V_{max}$) (**e**, *Ptchd1*$^{Y/+}$, 33.5 mV; *Ptchd1*$^{Y/-}$, 33.6 mV, p = 0.092; **n**, *Cul3*$^{+/+}$, 43.0 mV, *Cul3*$^{+/-}$, 41.9 mV, p = 0.984). (**f,o**), After-hyperpolarisation (AHP) amplitude (**f**, *Ptchd1*$^{Y/+}$, -63.3 mV; *Ptchd1*$^{Y/-}$, -60.7 mV, p < 0.001; **o**, *Cul3*$^{+/+}$, -63.8 mV, *Cul3*$^{+/-}$, -64.4 mV, p = 0.296). (**g,p**), Width at half-peak (**g**, *Ptchd1*$^{Y/+}$, 0.580 ms; *Ptchd1*$^{Y/-}$, 0.817 ms, p < 0.001; **p**, *Cul3*$^{+/+}$, 0.717 ms, *Cul3*$^{+/-}$, 0.736 ms, p = 0.565). (**h,q**), Rise time (**h**, *Ptchd1*$^{Y/+}$, 0.635 ms; *Ptchd1*$^{Y/-}$, 0.734 ms, p = 0.011; **q**, *Cul3*$^{+/+}$, 0.790 ms, *Cul3*$^{+/-}$, 0.784 ms, p = 0.399). (**i,r**), Exponential decay constant (**i**, *Ptchd1*$^{Y/+}$, 0.066; *Ptchd1*$^{Y/-}$, 0.043, p < 0.001; **r**, *Cul3*$^{+/+}$,0.052, *Cul3*$^{+/-}$, 0.067, p = 0.867). Points represent *Ptchd1*$^{Y/+}$ or *Cul3*$^{+/+}$ (grey) and *Ptchd1*$^{Y/-}$ (orange) or *Cul3*$^{+/-}$ (pink) dPAG cells. *Ptchd1*$^{Y/+}$, n = 4, (**a-d**) 16 cells, (**e-i**) 58 spikes; *Ptchd1*$^{Y/-}$, n = 4, (**a-d**) 16 cells, (**e-i**) 72 spikes. *Cul3*$^{+/+}$, n = 3, (**j-m**) 12 cells, (**n-r**) 68 spikes; *Cul3*$^{+/-}$, n = 3, (**j-m**) 12 cells, (**n-r**) 73 spikes. **s**, Summary of the relationship between current-injection and action potential firing showing a strong reduction in firing in *Ptchd1*$^{y/-}$ dPAG cells. Effect of genotype without **α**-DTX: P = 0.046; effect of **α**-DTX on *Ptchd1*$^{Y/-}$ firing: P = 0.680. (*Ptchd1*$^{Y/+}$, n = 4, 16 cells; *Ptchd1*$^{Y/-}$, n = 4, 20 cells; *Ptchd1*$^{Y/-}$ with **α**-DTX, n = 3, 13 cells). Inset, representative example traces to a 50 pA current injection for *Ptchd1*$^{Y/+}$ (black, top) and *Ptchd1*$^{Y/-}$ (orange, bottom). **t**, Average shape (left) and phase plane analysis (right) of the action potentials generated in the rheobase sweep (*Ptchd1*$^{Y/+}$, 16 cells, 58 spikes; *Ptchd1*$^{Y/-}$, 20 cells, 72 spikes). **u**, Summary of the relationship between current-injection and action potential firing in *Cul3* dPAG cells (*Cul3*$^{+/+}$, grey, 12 cells, n = 3; *Cul3*$^{+/-}$, pink, 12 cells, n = 3, P = 0.683 for the effect of genotype). **v**, Average shape (left) and phase plane analysis (right) of the action potentials generated in the rheobase sweep for *Cul3*$^{+/+}$ (grey, 68 spikes) and *Cul3*$^{+/-}$ (pink, 73 spikes). Box-and-whisker plots show median, IQR and range. Shaded areas represent SEM. Lines are shaded areas, mean ± s.e.m., respectively. p-values are Wilcoxon's rank-sum tests and P-values are two-way repeated measures ANOVA, unless specified.
